# Electrophysiological foundations of the human default-mode network revealed by brain-wide intracranial-EEG recordings during resting-state and cognition

**DOI:** 10.1101/2020.07.24.220566

**Authors:** Anup Das, Carlo de los Angeles, Vinod Menon

## Abstract

Investigations using noninvasive functional magnetic resonance imaging (fMRI) have provided significant insights into the unique functional organization and profound importance of the human default mode network (DMN), yet these methods are limited in their ability to resolve network dynamics across multiple timescales. Electrophysiological techniques are critical to address these challenges, yet few studies have explored the neurophysiological underpinnings of the DMN. Here we investigate the brain-wide electrophysiological organization of the DMN in a common large-scale network framework consistent with prior fMRI studies. We used brain-wide intracranial EEG (iEEG) recordings, and evaluated intra- and cross-network interactions during the resting-state and cognition. Our analysis revealed significantly greater intra-DMN phase iEEG synchronization in the slow-wave (< 4 Hz) while DMN interactions with other brain networks was higher in all higher frequencies. Crucially, slow-wave intra-DMN synchronization was observed in the task-free resting-state and during verbal memory encoding and recall. Compared to resting-state, intra-DMN phase synchronization was significantly higher during both memory encoding and recall. Slow-wave intra-DMN phase synchronization increased during successful memory retrieval, highlighting its behavioral relevance. Finally, analysis of nonlinear dynamic causal interactions revealed that the DMN is a causal outflow network during both memory encoding and recall. Our findings identify dynamic spectro-temporal network features that allow the DMN to maintain a balance between stability and flexibility, intrinsically and during task-based cognition, provide novel insights into the neurophysiological foundations of the human DMN, and elucidate network mechanisms by which it supports cognition.

---

The default mode network (DMN) is a large-scale distributed brain network which plays a critical role in cognition, including episodic memory formation and monitoring internal thoughts (Greicius MD et al. 2003; Buckner RL et al. 2008; Raichle ME 2015). DMN impairments are prominent in psychiatric disorders and this network is particularly sensitive to Alzheimer’s disease pathology and the ensuing loss of episodic memory and related cognitive functions (Greicius MD et al. 2004; Sheline YI et al. 2010; Staffaroni AM et al. 2018). In the past two decades, investigations using noninvasive functional magnetic resonance imaging (fMRI) techniques have provided significant insights into the unique functional organization of the human DMN. However, the electrophysiological basis and network properties of the DMN are poorly understood as fMRI does not have the requisite temporal resolution to address foundational questions in human systems neuroscience: what binds the DMN together as a network and segregates it from other large-scale brain networks? Here, for the first time, we address this question using brain-wide depth intracranial EEG (iEEG) recordings and investigate the neurophysiological foundations of the DMN, and its dynamic spectro-temporal properties in the resting-state and task-based cognition.

The default mode network (DMN) is a large-scale distributed brain network which plays a critical role in cognition, including episodic memory formation and monitoring internal thoughts (Greicius MD et al. 2003; Buckner RL et al. 2008; Raichle ME 2015). DMN impairments are prominent in psychiatric disorders and this network is particularly sensitive to Alzheimer’s disease pathology and the ensuing loss of episodic memory and related cognitive functions (Greicius MD et al. 2004; Sheline YI et al. 2010; Staffaroni AM et al. 2018). In the past two decades, investigations using noninvasive functional magnetic resonance imaging (fMRI) techniques have provided significant insights into the unique functional organization of the human DMN. However, the electrophysiological basis and network properties of the DMN are poorly understood as fMRI does not have the requisite temporal resolution to address foundational questions in human systems neuroscience: what binds the DMN together as a network and segregates it from other large-scale brain networks? Here, for the first time, we address this question using brain-wide depth intracranial EEG (iEEG) recordings and investigate the neurophysiological foundations of the DMN, and its dynamic spectro-temporal properties in the resting-state and task-based cognition.

The DMN consists of a distributed set of brain regions, including the posterior cingulate cortex, medial prefrontal cortex, angular gyrus, anterior prefrontal cortex, lateral temporal cortex and medial temporal lobe (MTL). Investigations of the electrophysiological properties of the DMN have been hampered by challenges of obtaining high-quality iEEG data from large-scale brain networks spanning widely distributed nodes across multiple lobes (Menon V et al. 1996; Freeman WJ et al. 2009; Parvizi J and S Kastner 2018). Furthermore, few electrophysiological investigations of the DMN have examined cortical recordings simultaneously with the MTL, an important constituent node of the DMN (Greicius MD *et al.* 2003). Here we overcome these limitations using brain-wide depth iEEG recordings spanning cortical and MTL nodes that constitute the DMN. Critically, the extensive distribution of electrodes allowed us to probe the spectro-temporal organization of the DMN in relation to six other large-scale cortical networks that have been consistently identified using fMRI: dorsal attention, ventral attention, frontoparietal, visual, motor, and limbic (Yeo BT et al. 2011). This approach allowed us to address critical questions regarding the electrophysiological properties of the DMN in relation to fMRI-derived functional architectures of the human brain.

Previous iEEG studies focusing on individual nodes of the DMN have demonstrated suppression of the posterior cingulate cortex during mental arithmetic (Dastjerdi M et al. 2011; Daitch AL and J Parvizi 2018), and activation of the retrosplenial cortex during autobiographical memory retrieval (Foster BL et al. 2013). Furthermore, transient suppression of gamma band activity is correlated with task complexity and performance on a visual search task (Ossandon T et al. 2011). Investigations of iEEG studies have also reported resting-state correlations in high-frequency band power fluctuations between the posterior cingulate cortex and angular gyrus (Foster BL et al. 2015), and between the posterior cingulate cortex and medial prefrontal cortex (Kucyi A et al. 2018) nodes of the DMN. However, other studies have failed to find such an association (Hacker CD et al. 2017), likely reflecting the small sample sizes and limited electrode coverage in prior studies (Fox KCR et al. 2018). Enhanced correlations in the theta frequency band have also been reported between lateral temporal cortex nodes of the DMN and the fronto-parietal central executive network (Kam JWY et al. 2019) as well as transient phase locking of the retrosplenial cortex with the medial temporal lobe during autobiographical memory retrieval (Foster BL *et al.* 2013). In other related research, resting-state correlations in the ultralow (<0.5 Hz) frequency have been reported in surface electrocorticogram recordings over somatosensory and motor cortex (He BJ et al. 2008), but it is not known whether such a frequency-specific network coupling also extends to the large-scale functional organization of the DMN. The limited electrode coverage in extant iEEG studies (please see **Table S1** for a summary of the most relevant studies) has made it impossible to examine connectivity across the distributed nodes that comprise the DMN and identify neurophysiological properties that segregate it from other large-scale brain networks, as identified using whole-brain fMRI.

A different line of research using scalp EEG recordings with concurrent fMRI has also probed the relation between fMRI activity and band-limited EEG fluctuations, but no consensus has emerged about the spectro-temporal organization of the DMN as the frequency-specificity of the correlations has been inconsistent across studies (Laufs H et al. 2003; Moosmann M et al. 2003; Mantini D et al. 2007; Scheeringa R et al. 2008; Jann K et al. 2009; Wu L et al. 2010; Yuan H et al. 2010). Crucially, the use of scalp EEG to characterize intra- and inter-network synchronization of the DMN is highly problematic due to volume conduction (Freeman WJ *et al.* 2009) and the inability of scalp EEG recordings to capture localized subcortical DMN regions such as the hippocampus.

Furthermore, little is known about how the large-scale intrinsic spectro-temporal organization and dynamic causal interactions of the DMN is modulated by cognition. Episodic memory is thought to be an important function associated with the DMN (Greicius MD *et al.* 2003; Greicius MD *et al.* 2004; Buckner RL *et al.* 2008; Raichle ME 2015), and iEEG recordings from individual brain regions have reported theta-band activity in the hippocampus and paraphippocampal gyrus during recall of verbal, temporal and spatial information from recently encoded memories (Watrous AJ et al. 2013; Jacobs J et al. 2016; Goyal A et al. 2018). However, the large-scale electrophysiological organization of the DMN and causal network dynamics during memory encoding and retrieval are not known and, crucially, no intracranial studies have probed how the intrinsic organization of the DMN and its interaction with other large-scale brain networks is altered by task-related episodic memory processes. More generally, there is growing evidence from fMRI studies that the DMN has a direct role in cognition across multiple cognitive domains, as revealed by task-related modulation of its posterior cingulate cortex, angular gyrus and middle temporal gyrus nodes (Greicius MD and V Menon 2004; Crittenden BM et al. 2015; Murphy C et al. 2018; Smith V et al. 2018; Sormaz M et al. 2018; Murphy C et al. 2019). A deeper understanding of the role of the DMN during cognition requires clarification of the electrophysiological mechanisms that support task-related network interactions in relation to its intrinsic spectro-temporal organization.

A systematic analysis using depth iEEG recordings spanning distributed nodes of the DMN, including the PCC, MTL, angular gyrus, and anterior temporal cortex, in parallel with other large-scale brain networks is needed to resolve these discrepancies and address fundamental questions regarding the integrity of this critical brain network (Hacker CD *et al.* 2017). To address this challenge, we used iEEG data from the UPENN-RAM study (Solomon EA et al. 2019) that included brain-wide depth recordings from 102 participants and 12,780 electrodes, from which we selected 36 participants with 879 electrodes spanning seven fMRI-derived brain networks (Yeo BT *et al.* 2011). This allowed us to probe intra-DMN connectivity and contrast it with DMN interactions with other brain networks in a common large-scale network framework consistent with prior whole-brain non-invasive imaging studies.

The first main goal of our study was to probe the intrinsic spectro-temporal organization of the human DMN during a task-free “resting” state and determine frequency-specific instantaneous phase synchronization measures that capture linear as well as nonlinear intermittent and nonstationary dynamics observed in iEEG data (Menon V *et al.* 1996; Lachaux JP et al. 1999). We hypothesized that intra-DMN network interactions would be significantly stronger than inter-network interactions in the low frequencies (< 4 Hz), based on the hypothesized role of the ultralow (< 0.5 Hz) and delta (0.5-4 Hz) frequency bands in large-scale network synchronization (Buzsáki G 2000). The second major goal of our study was to extend our analysis and methodology to investigate DMN phase synchronization during an episodic memory task involving encoding and recall of a list of words. Investigations of intra- and cross-network interactions during verbal episodic memory is particularly relevant in the context of DMN function since the PCC, MTL, angular gyrus, and anterior temporal cortex have each been consistently implicated in verbal episodic and semantic memory (Binder JR and RH Desai 2011; Hasson U et al. 2015). The final goal of our study was to determine how DMN synchrony changes during cognition, when compared to rest. We hypothesized that intra-DMN network synchronization would increase during memory encoding and recall when compared to the resting-state while preserving the overall low-frequency-dependent synchrony of the DMN. Our study provides novel, behaviorally and functionally relevant, insights into the neurophysiological foundations of the human DMN and its role in cognition.

## Results

### Slow-wave intra-DMN synchronization during the resting-state

We first examined intrinsic intra-network synchronization of the DMN and contrasted it with cross-network synchronization with six other brain networks, identified using the fMRI-based cortical network atlas (Yeo BT *et al.* 2011) (**Figures 1a, S1, S2, Tables S2–S5; Methods**). iEEG data across the non-DMN networks were combined to limit multiple comparison testing, and more directly address our specific hypotheses. Analysis of instantaneous phase locking values (PLVs) revealed significantly greater intra-DMN phase synchronization, compared to DMN interactions with the other six networks, in the ultralow-delta band (< 4 Hz) (*p*<0.001, Mann-Whitney U-test, FDR-corrected for multiple comparisons) (**Figure 2a**). This pattern was reversed in the higher frequency bands, with stronger cross-network, compared to intra-DMN, synchronization in the theta, alpha, beta, and gamma bands (all *ps*<0.001, Mann-Whitney U-test, FDR-corrected) (**Figure 2a**). Note the gradual decrease in the PLV values from low to high frequency bands, consistent with the well-known 1/f spectral scaling in EEGs (He BJ et al. 2010).

**Figure 1.**
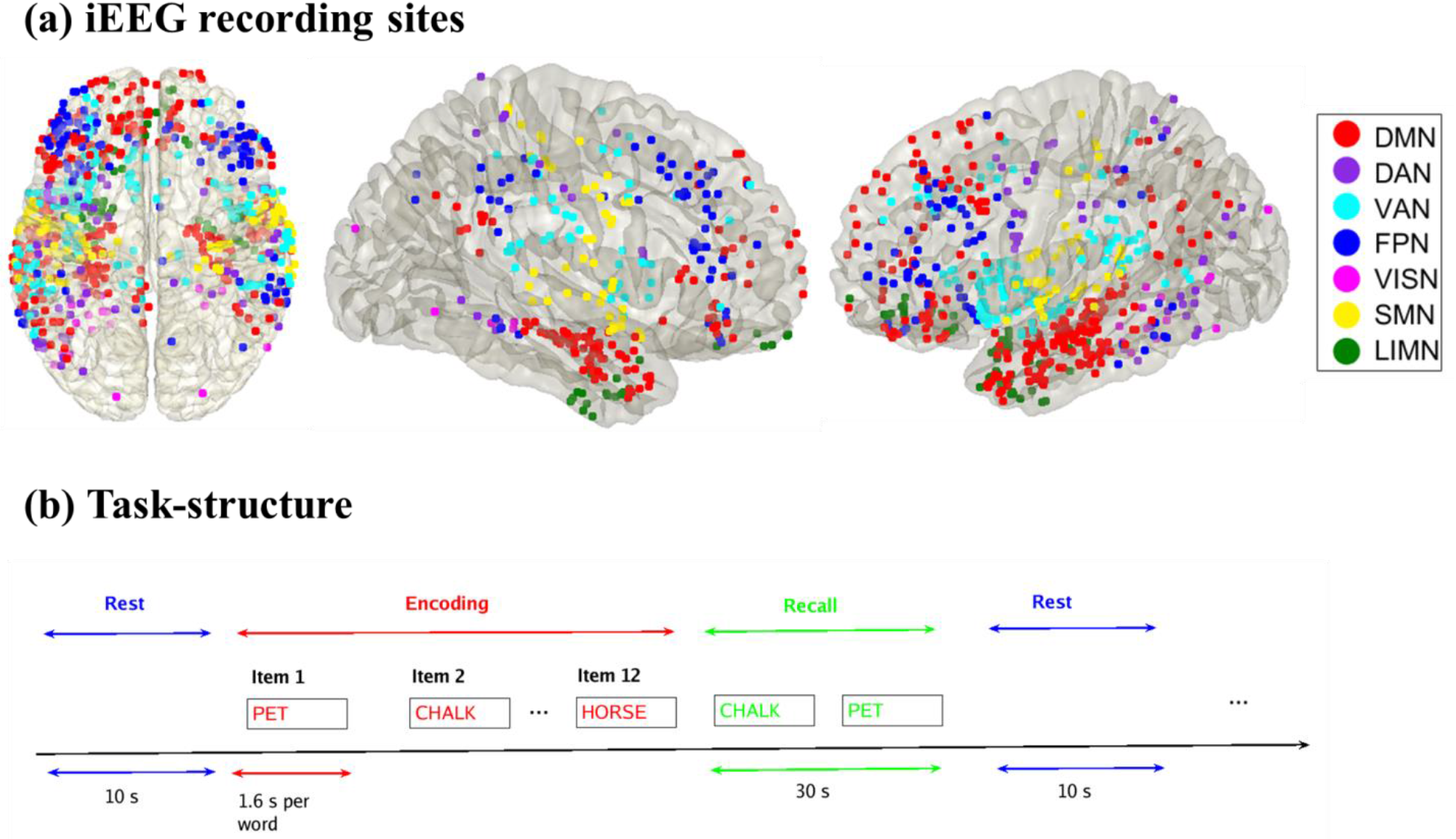
**(a) iEEG recording sites for the 7 fMRI-derived brain networks investigated in this study.** The Yeo cortical atlas was used to map the default mode (DMN), dorsal attention (DAN), ventral attention (VAN), frontoparietal (FPN), visual (VISN), somatomotor (SMN), and limbic (LIMN) networks. In addition to cortical areas, the DMN also included hippocampal regions determined using the Brainnetome atlas (**Figures S1, S2). (b) Cognitive task structure.** Participants performed multiple trials of a “free recall” experiment, where they were first presented with a list of words and later asked to recall as many as possible from the original list (see **Methods** for details).

**Figure 2.**
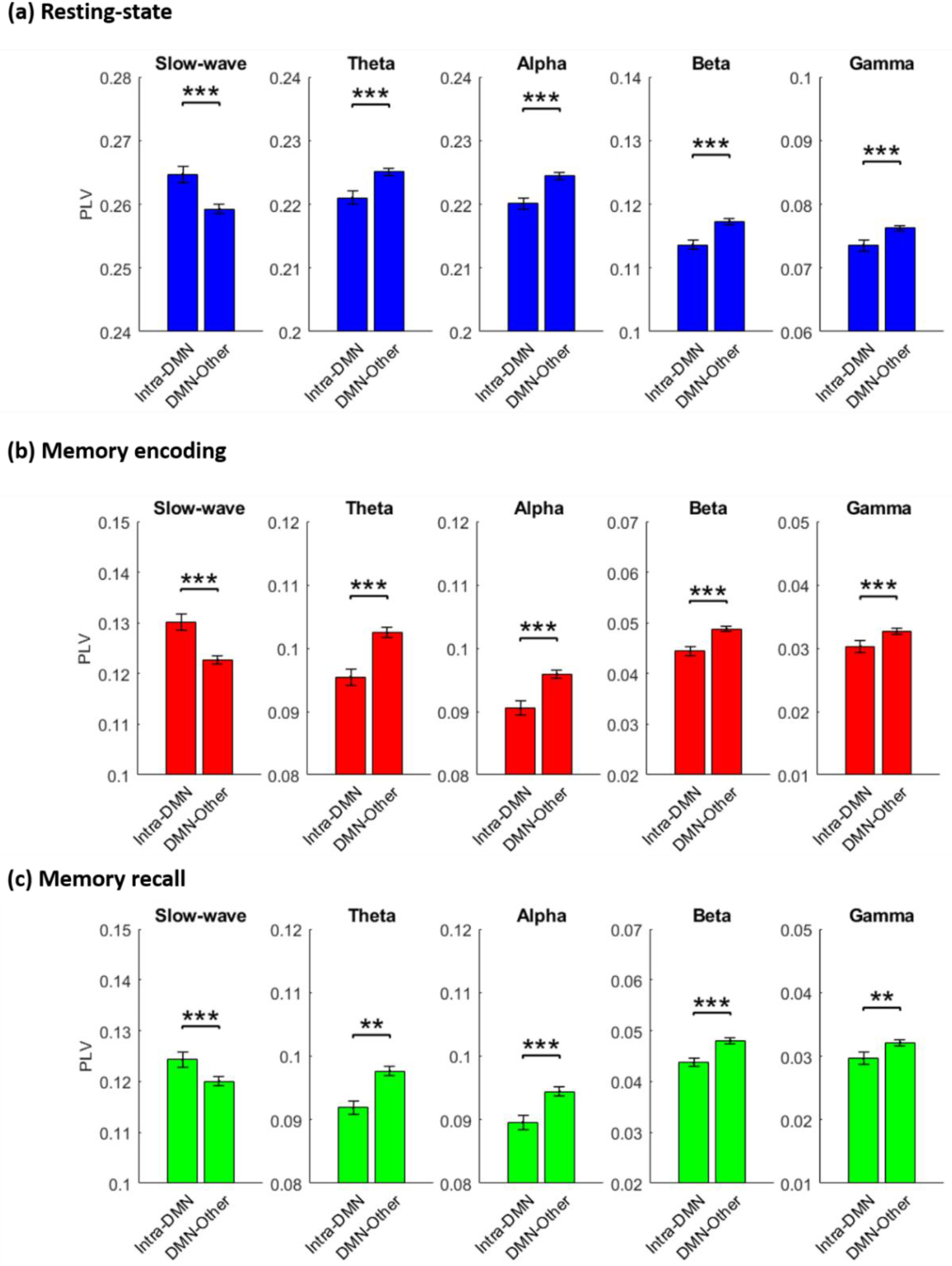
**Intra-DMN phase synchronization during (a) Resting-state, (b) Memory encoding, and (c) Memory recall.** In all three conditions, intra-DMN connectivity was characterized by synchronization in the slow-wave frequency band (< 4 Hz) while cross-network interaction of the DMN was dominated by higher frequencies. Intra-DMN denotes phase locking values (PLVs) between DMN electrodes and DMN-Other denotes PLV between DMN electrodes and electrodes in the 6 other brain networks. Error bars denote standard error of the mean (SEM) across all pairs of electrodes. \*\*\**p* < 0.001, ** *p* < 0.01 (FDR-corrected *q*<0.05).

These results demonstrate that intra-DMN connectivity is dominated by ultralow-delta (hereafter referred to as slow-wave) synchronization, and that cross-network interaction of the DMN is dominated by higher frequencies, thus providing novel evidence for spectral segregation of the DMN in the resting-state.

### Slow-wave intra-DMN synchronization during memory encoding

We then extended the above analysis to the memory encoding period of a verbal episodic memory task in which participants were presented with a sequence of words and asked to remember them for subsequent recall (**Figure 1b, Methods**). Again, intra-DMN slow-wave phase synchronization was higher when compared to DMN interactions with the other 6 networks (*p*<0.001, Mann-Whitney U-test, FDR-corrected) (**Figure 2b**). The reverse was true in higher frequency bands: DMN electrodes had higher phase synchronization with electrodes outside the DMN than intra-DMN electrodes in the theta, alpha, beta, and gamma bands (all *ps*<0.001, Mann-Whitney U-test, FDR-corrected) (**Figure 2b**).

These results demonstrate that intra-DMN connectivity is dominated by slow-wave synchronization, and that cross-network interaction of the DMN is dominated by higher frequencies, thus providing evidence that spectral segregation of the DMN, observed intrinsically, is also maintained during memory encoding.

### Slow-wave intra-DMN synchronization during memory recall

Next, we examined phase synchronization during the recall phase of the verbal episodic memory task in which participants recalled the words they had seen during the memory encoding phase. Here again, intra-DMN slow-wave phase synchronization was higher compared to DMN interactions with the other 6 networks (*p*< 0.001, Mann-Whitney U-test, FDR-corrected) (**Figure 2c**). The reverse was again true in higher frequency bands: DMN electrodes had higher phase synchronization with electrodes outside the DMN than intra-DMN electrodes in the theta (*p*<0.01), alpha (*p*<0.001), beta (*p*<0.001), and gamma (*p*<0.01) bands (Mann-Whitney U-Test, FDR-corrected) (**Figure 2c**). For control analyses related to amplitude fluctuations in high-gamma band (80-160 Hz), see **Supplementary Results**.

These results demonstrate that intra-DMN connectivity is dominated by slow-wave synchronization, and that cross-network interaction of the DMN is dominated by higher frequencies, thus providing evidence that spectral segregation of the DMN, observed intrinsically, is also maintained during memory recall.

### Intra- and cross-network phase synchronization of the DMN sans hippocampus

Most human functional neuroimaging research on the DMN and fMRI-based atlases of brain networks have focused on its cortical nodes. We investigated the extent to which spectral segregation of the DMN was dependent on the hippocampus. We conducted additional analyses probing intra- and cross-network phase synchronization during resting-state, memory encoding, and memory recall conditions excluding hippocampus electrodes from the DMN. This analysis revealed that intra-DMN slow-wave phase synchronization was higher compared to DMN interactions with the other 6 networks in all three conditions (*ps*<0.001, Mann-Whitney U-test, FDR-corrected) (**Figure S3**). Consistent cross-network interaction of the DMN was observed in all three task conditions in the beta band (*ps*<0.001, Mann-Whitney U-test, FDR-corrected). These results suggest that key intra- and cross-network phase synchronization properties of the DMN hold independent of the hippocampus, but are strengthened by its inclusion.

### Broad-band increases in intra-DMN phase synchronization during memory encoding and memory recall, compared to resting state

We next investigated intra-DMN synchronization changes during memory encoding and recall, compared to the resting-state. The duration of task and rest epochs were matched to ensure that differences in network dynamics could not be explained by the differences in the duration of the epochs. Our analysis revealed that intra-network phase synchronization of the DMN during both the encoding and recall phases of the verbal episodic memory task was higher than that during rest (*ps*<0.001 in all frequency bands (slow-wave, theta, alpha, beta, and gamma) during both the encoding and recall task conditions, except one case where *p*<0.01, Mann-Whitney U-test, FDR-corrected) (**Figure 3**). These results demonstrate broad-band increases in phase synchronization within the DMN during both memory encoding and recall.

**Figure 3.**
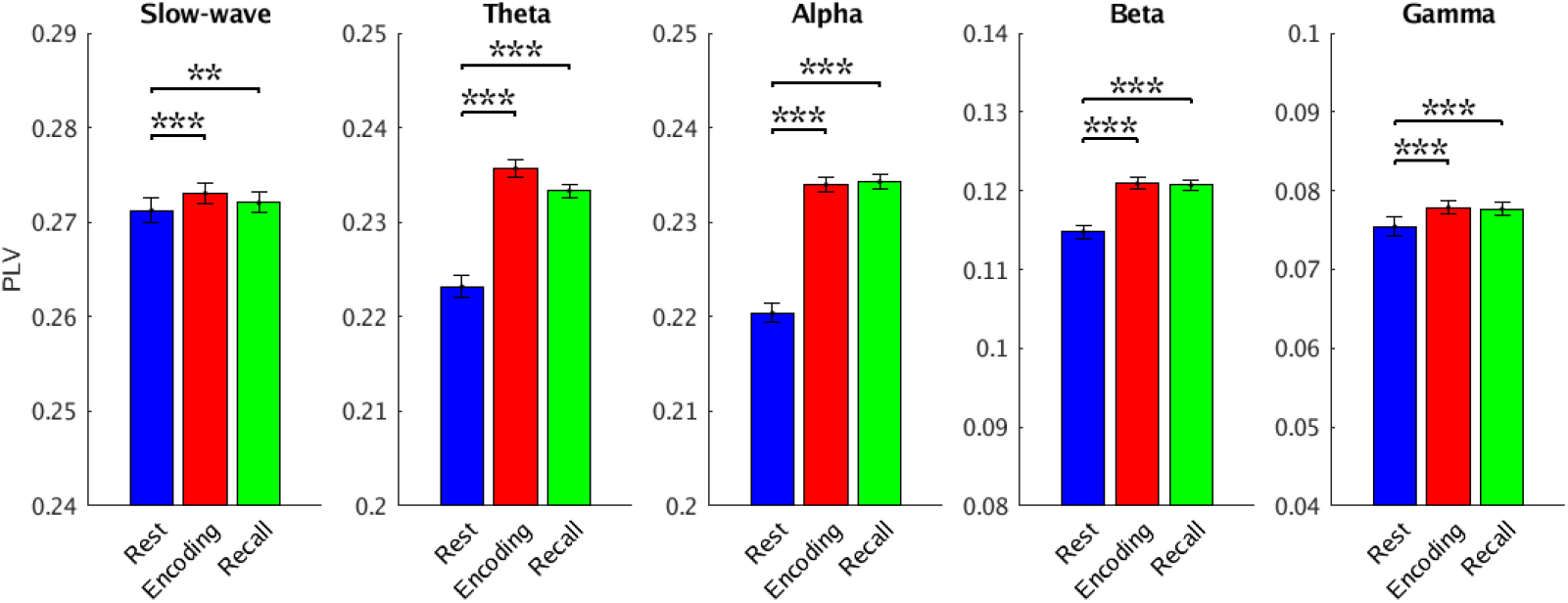
**Intra-DMN phase synchronization during memory encoding and memory recall, compared to resting-state.** Intra-DMN phase synchronization, assessed using phase locking values (PLVs), was higher during both memory encoding and recall compared to resting-state, in the slow-wave and all other frequency bands. Error bars denote standard error of the mean (SEM) across all pairs of electrodes. \*\*\**p* < 0.001, \*\**p* < 0.01 (FDR-corrected *q*<0.05).

### Increased slow-wave intra-DMN phase synchronization associated with successful vs. unsuccessful memory recall

Next, we sought to investigate the behavioral significance of intra-DMN slow-wave phase synchronization. We examined whether intra-DMN phase synchronization differed between trials that were subsequently recalled correctly versus those that were not. During the encoding phase of the memory task, intra-DMN slow-wave phase synchronization was higher for successfully, versus unsuccessfully, recalled words (*p*<0.001, Wilcoxon signed-rank test) (**Figure 4a**). This finding was replicated in the memory recall phase of the task (*p*<0.001, Wilcoxon signed-rank test) (**Figure 4b**). These results suggest that spectral integrity of the DMN is associated with successful memory performance.

**Figure 4.**
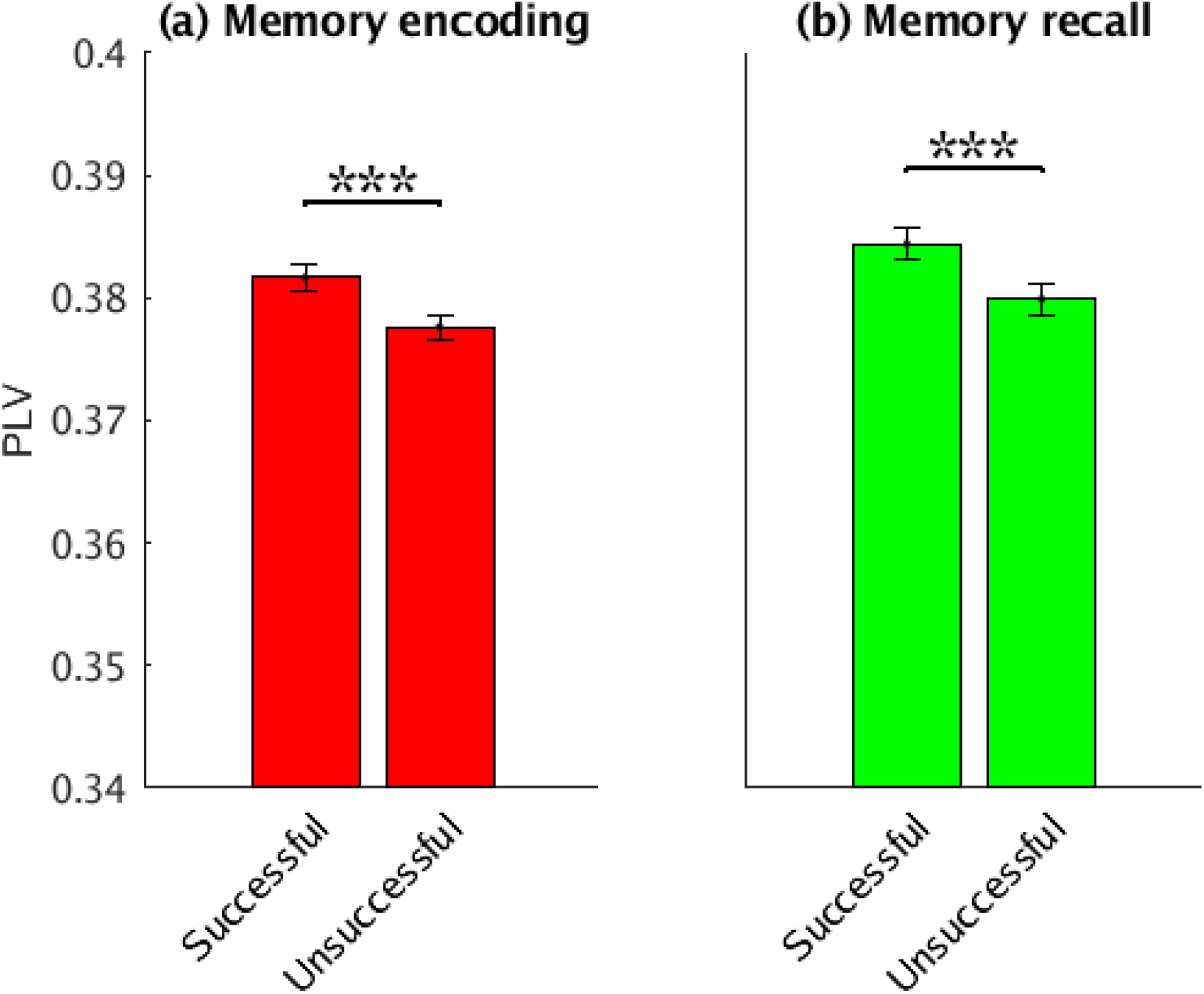
**Intra-DMN phase synchronization during (a) successful vs. unsuccessful Memory encoding and (b) successful vs. unsuccessful Memory recall.** Intra-DMN phase synchronization was higher during successful encoding/recall compared to unsuccessful encoding/recall in the slow-wave frequency band. Error bars denote standard error of the mean (SEM) across all pairs of electrodes. \*\*\**p* < 0.001.

### Enhanced causal influences from the DMN to other networks during memory encoding and memory recall

To investigate the role of dynamic causal interactions of the DMN in episodic memory task performance, we used PTE (Lobier M et al. 2014) to compute the net causal outflow from the DMN to the other six networks (see **Methods** for details). During both the memory encoding and recall phases, the DMN showed greater net causal outflow to other networks than the reverse (*ps*<0.001, Wilcoxon signed-rank test) (**Figures 5a, b**). These results demonstrate that the DMN plays an important causal role in interactions with other networks during episodic memory formation.

**Figure 5.**
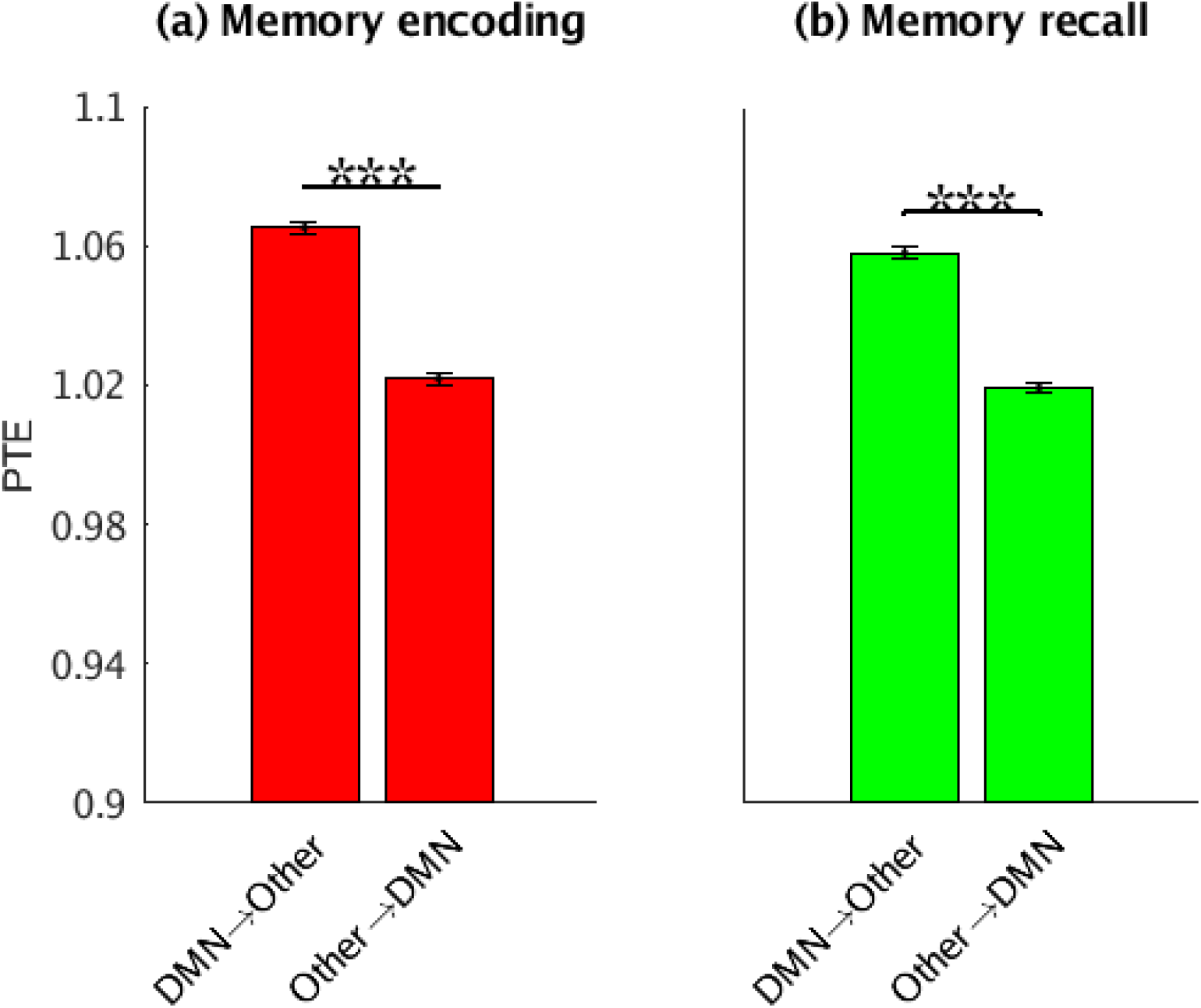
**Causal network influences, measured using phase transfer entropy (PTE), during (a) Memory encoding and (b) Memory recall.** The DMN showed significantly higher net causal outflow to the 6 other networks, than the reverse. DMN→Other denotes PTE from DMN electrodes to electrodes in the 6 other brain networks; Other→DMN denotes PTE from electrodes in the 6 other networks to the DMN. Error bars denote standard error of the mean (SEM) across all pairs of electrodes. \*\*\**p* < 0.001.

Finally, comparison of causal outflow from the DMN to other networks across conditions revealed greater causal outflow during the resting-state compared to both memory encoding and recall (*ps*<0.01, Mann-Whitney U-test, FDR-corrected) (**Figure 6**). Causal outflow from the DMN was greater during memory encoding compared to recall (*p*<0.01, Mann-Whitney U-test, FDR-corrected) (**Figure 6)**. These results suggest that the DMN also exerts strong causal influences on other brain networks intrinsically and is not specific to task-related memory processes.

**Figure 6.**
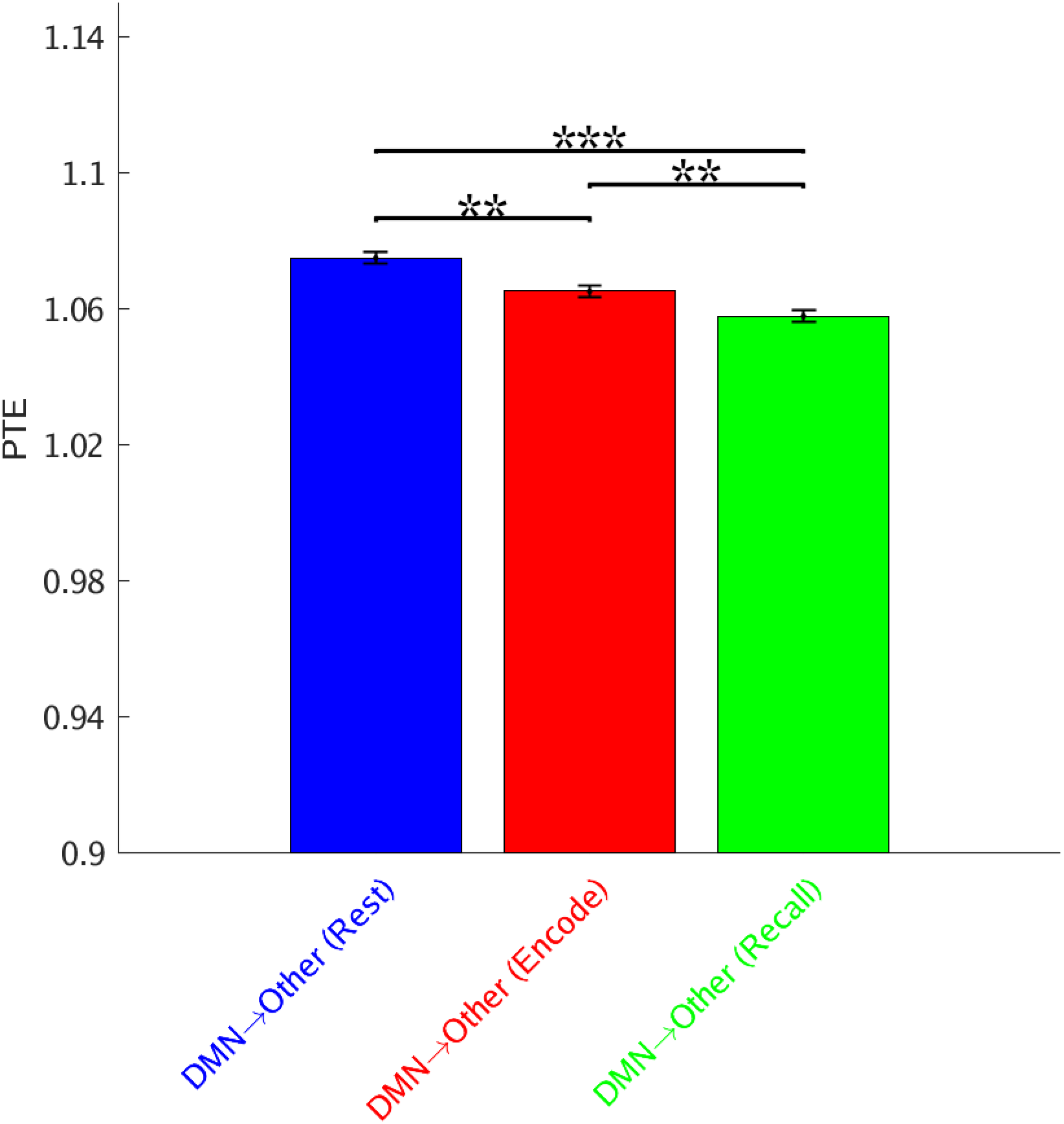
**Causal network influences of the DMN on other networks during resting-state, memory encoding, and memory recall.** Causal outflow from the DMN was higher during resting-state compared to both the memory encoding and memory recall. Causal outflow from the DMN was also higher during memory encoding compared to memory recall. Error bars denote standard error of the mean (SEM) across all pairs of electrodes. *** *p* < 0.001, ** *p* < 0.01 (FDR-corrected *q*<0.05).

### DMN power spectral density does not differ from other networks

To determine whether synchronization within the DMN was driven by differences in the amplitude of iEEG fluctuations, we compared power spectral density (see **Methods** for details) in the DMN and the other six large-scale networks. This analysis revealed that power in the DMN did not differ from power in the other six networks in any of the frequency bands during all three task conditions (all *ps*>0.05, Mann-Whitney U-test). These results suggest that synchronization within the DMN is not driven by differences in the amplitude of iEEG fluctuations (see **Supplementary Results** for further details).

### Elevated slow-wave intra-DMN phase synchronization is not related to inter-electrode distance

We performed additional control analysis to rule distance out as a potential factor for the higher intra-DMN phase synchronization in the slow-wave frequency band. We calculated the Euclidean distance between all pairs of intra-DMN electrodes and for all pairs with one electrode in the DMN and the second in one of the six other networks. The distance of intra-DMN electrodes was, in fact, higher than the cross-DMN electrodes distance (*p*<0.001, Mann-Whitney U-test) (**Figure S4**). To further rule out the possibility that low-frequency synchronization may arise from greater inter-electrode distance within the DMN, we examined the relation between PLV and distance in resting-state iEEG. We found no significant correlation between the two (*r* = −0.02, *p* > 0.27, Pearson) (**Figure S5**). These results suggest that the intra-DMN spectral integration in the slow-wave frequency band is a reflection of the strong instantaneous phase coupling among the intra-DMN brain regions as captured by the phase locking values.

### Intra- vs. cross-network synchronization of the six other networks

Finally, we conducted additional control analyses of intra- vs. cross-network phase synchronization of the other 6 networks: dorsal attention, ventral attention, frontoparietal, visual, motor, and limbic (**Figures S6–S11** respectively). Dorsal attention, ventral attention, visual, somatomotor, and limbic networks did not show consistent intra-network phase synchronization in the slow-wave frequency band across the resting-state, memory encoding, and memory recall conditions *(ps* > 0.05, Mann-Whitney U-test, FDR-corrected) (**Figures S6–S7, S9–S11**). The frontoparietal network showed intra-network phase synchronization in slow-wave frequency band across the resting-state, memory encoding, and memory recall conditions *(ps* < 0.01, Mann-Whitney U-test, FDR-corrected) (**Figure S8**). However, the frontoparietal network did not show consistent cross-network signatures in the higher frequency bands across the three conditions, unlike the DMN *(ps* > 0.05, Mann-Whitney U-test, FDR-corrected) (**Figure S8**).

These control analyses suggest that the pattern of stronger intra-network phase synchronization in the slow-wave frequency band and increased cross-network synchronization in the theta, alpha, beta, and gamma frequency bands across resting-state, memory encoding, and memory recall conditions is not consistently observed in the other six large-scale brain networks (**Table S6**).

## Discussion

Using brain-wide electrophysiological recordings spanning seven large-scale brain networks, we addressed the following key questions to probe DMN function in a common network framework with fMRI studies: (1) What are the foundational electrophysiological properties of the human DMN which allows it to function as a coherent network? (2) What are the mechanisms underlying synchronization and causal interactions of the DMN with other networks during stimulus-driven cognition? To probe the spectro-temporal dynamics of the DMN, we examined phase locking and transfer entropy measures which are better suited to capturing nonlinear and nonstationary synchronization as well as causal dynamics associated with intra- and cross-network interactions (Menon V *et al.* 1996; Lachaux JP *et al.* 1999; Bruns A 2004; Lopour BA et al. 2013; Lobier M *et al.* 2014; Hillebrand A, P Tewarie, E van Dellen, et al. 2016). We found that intra-DMN connectivity is dominated by slow-wave (< 4 Hz) synchronization, relative to DMN interactions with other brain networks. This pattern was observed intrinsically and during both the encoding and recall phases of a verbal episodic memory task. DMN synchronization increased in all frequency bands during memory processing, but this increase did not alter slow-wave integration and higher-frequency segregation of the DMN from the other six large-scale cortical networks. Our findings provide novel evidence for frequency-specific segregation and integration of the DMN from other brain networks during rest and its maintenance during cognition. More generally, our findings advance knowledge of the neurophysiological foundations of the DMN, and clarify dynamic neural mechanisms underlying its role in task-based cognition.

### Intrinsic spectro-temporal network organization of the DMN

The first goal of our study was to characterize the intrinsic spectro-temporal organization of the DMN. Analysis of the spectral properties of the human DMN has been necessarily constrained by limited placement of iEEG electrodes, consequently previous work has mainly focused on activation and deactivation in isolated nodes of the network (Ossandon T *et al.* 2011; Lopour BA *et al.* 2013; Foster BL *et al.* 2015; Hacker CD *et al.* 2017; Daitch AL and J Parvizi 2018; Solomon EA *et al.* 2019). It is currently impossible to acquire iEEG data across the entire human brain with resolutions matching those of fMRI. This limitation motivated our approach of using the network atlas of Yeo et al. (Yeo BT *et al.* 2011), which allowed us to fill critical gaps in our knowledge of the neurophysiological underpinnings of the human DMN vis-a-vis the dorsal attention, ventral attention, frontoparietal, visual, motor, and limbic networks. Importantly, for the purposes of the present study, this 7-network atlas was coarse-grained enough to allow sampling of a large number of participants who had simultaneous iEEG recordings spanning the DMN as well as the six other cortical networks. Furthermore, such atlases have been widely used to investigate resting-state and task-based interactions using fMRI (Menon V 2015). Note that in contrast to previous iEEG studies (Dastjerdi M *et al.* 2011; Foster BL *et al.* 2015; Daitch AL et al. 2016; Daitch AL and J Parvizi 2018; Kucyi A *et al.* 2018; Raccah O et al. 2018; Kucyi A et al. 2020), we took an unbiased approach for assigning electrodes to individual brain networks and we did not select electrodes based on arbitrary task activation or deactivation profiles. Our approach thus allowed us to more directly probe the electrophysiological foundations of the DMN as identified in fMRI studies, in line with our study goals. Crucially, compared to subdural recordings, bipolar depth recordings in the UPENN-RAM dataset provide a more effective way to capture local iEEG signals (Ekstrom AD and AJ Watrous 2014; Jacobs J *et al.* 2016; Goyal A *et al.* 2018). These advances allowed us to probe intra- and cross-network dynamics of the DMN in a common framework with fMRI studies.

We evaluated inter-regional synchronization of iEEG time series using phase-locking values (PLVs), based on the notion that two connected brain areas generate signals whose instantaneous phases evolve together (Lopour BA *et al.* 2013; Lobier M *et al.* 2014). We contrasted intra- and cross-network phase synchronization of the DMN and found greater PLV among DMN electrodes in a combined slow-wave frequency band that encompasses the delta (0.5-4 Hz) and a lower ultralow (< 0.5 Hz) band. Control analysis revealed that this elevated slow-wave intra-DMN phase synchronization was not related to inter-electrode distance. As the length of each verbal memory trial was ~4 s (see **Methods**) which did not allow us to analyze the ultralow band separately, we combined these frequency bands into a single ultralow-delta (< 4 Hz) ‘slow-wave’ band (Dalal SS et al. 2011). Our findings are consistent with previous reports of synchronized delta frequency oscillations correlated with resting-state fMRI (Lu H et al. 2007), and with slow-wave resting-state correlations observed in electrocorticogram recordings over somatosensory and motor cortex (He BJ *et al.* 2008) and in iEEG recordings from the anterior insula and anterior cingulate cortex nodes of the salience network (Das A and V Menon 2020).

In contrast to this pattern of slow-wave dominated iEEG synchronization within the DMN, we uncovered a consistent pattern of cross-network interaction of the DMN which was dominated by higher frequencies ranging from the theta (4-8 Hz), alpha (8-12 Hz), beta (12-30 Hz), to gamma (30-80 Hz) bands. This pattern is consistent with high frequency desynchrony between hippocampus and prefrontal cortex reported in iEEG recordings (Solomon EA et al. 2017). Our findings are also consistent with theoretical models and neural network simulations of hippocampal-neocortical circuits hinting at low frequency (delta) synchronization and high frequency (alpha) desynchronization (Parish G et al. 2018). Based on these observations, we suggest that stronger intra-network phase synchronization in the slow-wave and stronger cross-network phase synchronization in higher frequency bands may enable the DMN to maintain a frequency-specific balance between stability and flexibility in its interactions with other large-scale brain networks.

### Phase synchronization, rather than spectral power density, separates the DMN from other large-scale networks

Phase locking values as used in the present study provide a robust measure of instantaneous synchronization between electrode pairs (Menon V *et al.* 1996; Lachaux JP *et al.* 1999; Lopour BA *et al.* 2013; Lobier M *et al.* 2014). Previous findings using multielectrode array recordings in both humans and animal models have established that phase, rather than amplitude, is responsible for both spatial and temporal encoding of information in the brain (Lachaux JP *et al.* 1999; Kayser C et al. 2009; Siegel M et al. 2009; Lopour BA *et al.* 2013; Ng BS et al. 2013). In line with this computation, we found no differences in overall power between the DMN and any of the other six cortical networks in any frequency band. This issue is pertinent because of the 1/f characteristic of iEEG, reflecting significantly greater power at lower frequencies which might have resulted in increased amplitude of fluctuations and signal-to-noise ratio within the DMN. Taken together, these results suggest that slow-wave phase synchronization within the DMN is not driven by differences in the amplitude of iEEG fluctuations. Rather, the spectro-temporal segregation of the DMN is a reflection of frequency-specific instantaneous phase as accurately captured by PLV measures.

### DMN phase synchronization and spectral integrity is preserved during memory encoding and recall

The next goal of our study was to investigate the spectral integrity of the DMN during cognition. Episodic memory is one of the putative functions ascribed to the DMN (Greicius MD *et al.* 2003; Greicius MD *et al.* 2004; Buckner RL *et al.* 2008). Previous studies and theoretical viewpoints have suggested that the DMN is integral for autobiographical memory, thinking about episodic events from the past, and planning the future (Buckner RL *et al.* 2008; Binder JR and RH Desai 2011). A related line of research has shown that the posterior cingulate cortex, angular gyrus, and anterior temporal lobe nodes of the DMN are also involved in semantic associations that modulate the formation of new episodic associations (Long NM and MJ Kahana 2017). Despite considerable evidence for the involvement of individual DMN nodes in memory formation and recall, to our knowledge, no studies have examined network-level neurophysiological organization of the DMN and its interaction with other large-scale brain networks during memory processing.

To address this, we first examined frequency-specific iEEG phase synchronization during a verbal episodic memory task in which participants had to subsequently recall a list of visually-presented words (Solomon EA *et al.* 2019). Our analysis procedures were identical to those used in the case of resting-state iEEG data. We found that intra-DMN slow-wave phase synchronization was significantly greater than DMN interactions with the six other large-scale networks. In contrast, the reverse was true with respect to cross-network interactions of the DMN, which showed greater synchronization of the DMN in the theta, alpha, beta, and gamma bands with other networks when compared to intra-DMN synchronization. We repeated this analysis with iEEG acquired during recall of the previously encoded words and uncovered the same pattern of frequency-specific synchronization: greater intra-DMN slow-wave synchronization and greater cross-network synchronization in all other higher frequencies. Furthermore, this pattern of frequency-specific synchronization was strengthened by the inclusion of the hippocampus, consistent with its hypothesized involvement in DMN function (Watrous AJ *et al.* 2013; Solomon EA et al. 2018; Solomon EA *et al.* 2019).

These results demonstrate that intra-DMN connectivity is dominated by slow-wave synchronization during both memory encoding and recall. Task-related connectivity patterns thus recapitulate and extend features observed with resting-state iEEG and points to a pattern of DMN organization that is stable under perturbations induced by episodic memory. In sum, our analysis demonstrates that the frequency-specific spectral integrity of the DMN and its segregation from other large-scale networks is maintained during cognition. We suggest that a unique combination of frequency-specific synchronization and desynchronization may facilitate formation and break-up of functionally relevant neuronal assemblies associated with the DMN (Ahn S and LL Rubchinsky 2013).

### DMN phase synchronization increases during cognition, compared to rest, and is behaviorally relevant

The final goal of our study was to determine how DMN synchrony changes during cognition, when compared to rest. To address this, we compared intra-DMN synchronization during memory processing and the resting-state using PLVs. We found that intra-DMN slow-wave synchronization was significantly higher during both memory encoding and recall, compared to the resting-state. Furthermore, increased synchronization during memory encoding and recall, compared to rest, was also observed in higher frequency bands ranging from the theta, to gamma bands. This pattern of broad-band spectral changes provides novel electrophysiological evidence that the DMN plays a critical role in episodic memory formation. Consistent with this view, analysis of phase transfer entropy revealed enhanced causal interactions from DMN to other large-scale networks during both memory encoding and recall.

The lack of concurrent brain-wide recordings has limited our ability to characterize frequency-specific synchronization associated with large-scale brain networks such as the DMN. Previous investigators focusing on regional response during verbal memory recall have reported increased iEEG activity in multiple frequency bands along with increased theta-band synchronization within the MTL (Long NM and MJ Kahana 2017; Solomon EA *et al.* 2019). In contrast, greater synchronization in the delta-band between MTL and inferior and middle frontal gyri electrocorticograms was reported during spatial memory recall (Watrous AJ *et al.* 2013; Ekstrom AD and AJ Watrous 2014; Neuner I et al. 2014). Extending these findings specifically to the DMN organization at a network-level, we found greater phase synchronization during both the encoding and retrieval phases of the verbal episodic memory task when compared to the resting-state.

Crucially, memory-related increase in DMN phase synchronization was behaviorally relevant. Phase synchronization was higher for words that were subsequently recalled correctly, and these increases were observed during both the memory encoding and recall phases of the task. Importantly, increased intra-DMN phase synchronization during successful encoding/recall compared to unsuccessful encoding/recall was not driven by differences in the amplitude of iEEG fluctuations. Memory success-related changes were observed only in the slow-wave spectral range, consistent with a previous report of frequency-specific network connectivity during spatial memory retrieval (Watrous AJ *et al.* 2013). Our findings are also consistent with reports of higher phase synchronization in the delta band associated with successful memory retrieval (Jacobs J et al. 2007; Fell J et al. 2008) and memory-related teleportation (Vass LK et al. 2016), although these recordings were solely confined to the hippocampus. Our analysis of the extent to which spectral segregation of the DMN was dependent on the hippocampus revealed that key intra- and cross-network phase synchronization properties of the DMN held without the inclusion of hippocampal electrodes, but were enhanced by its inclusion. In an advance over prior studies, frequency-specific increases in our analysis are based on iEEG recordings from the major cortical and MTL regions of the entire DMN providing novel evidence of network level signatures of successful memory retrieval in the human DMN. Findings support the view that network-wide synchronization is a key feature of memory formation and emphasize the role of the DMN in this process (Hebscher M and JL Voss 2020).

Our findings thus address an important gap in the literature and suggest that large-scale DMN phase synchronization during memory formation is not limited to the delta band nor is it specific to recall of spatial information. Notably, compared to the resting-state, synchronization increased in all frequency bands ranging from the ultralow to gamma bands. These increases were not related to overall iEEG amplitude, as spectral power during both memory tasks did not differ from rest. It is noteworthy that despite broad-band increases in cross-network synchronization, DMN integrity was still preserved during memory formation. Our results highlight the role of the DMN as a whole in memory encoding and retrieval, and further suggest that higher spectral integrity of this network is associated with memory-related cognitive functions.

### Causal dynamics of the DMN

Finally, we examined the directionality of information flow between the DMN and other networks as regions whose activities are not instantaneously synchronized may interact via time-delayed causal influences. Granger causal analyses of fMRI time series suggest that multiple DMN nodes, including the posterior cingulate cortex and the ventromedial prefrontal cortex may exert influence on lateral frontoparietal cortex (Uddin LQ et al. 2009) rather than the other way around. However, it is unclear whether these results are an artifact of regional variations in hemodynamic response in fMRI which confound Granger causal measures, or whether they truly reflect an underlying neuronal process. Critically, previous iEEG studies have not examined causal interaction of the DMN with other large-scale brain networks during cognition. To address this question, we used phase transfer entropy (PTE), which provides a robust and powerful measure for characterizing information flow between brain regions based on phase coupling (Lobier M *et al.* 2014; Hillebrand A, P Tewarie, Ev Dellen, et al. 2016; Wang MY et al. 2017). PTE has several advantages over Granger causal analysis (Barnett L and AK Seth 2011) and transfer entropy (Schreiber T 2000; Lobier M *et al.* 2014), as it can capture nonlinear interactions, is more accurate than transfer entropy, and it estimates causal interactions based on phase, rather than amplitude, coupling. Taking advantage of the 1,000 Hz sampling rate of the UPENN-RAM cohort iEEG data, we examined causal outflow from the DMN to the other six networks and contrasted it with inflow into the DMN from the other networks.

Our analysis revealed that the DMN showed significantly greater net causal outflow to other networks, rather than the reverse, during both memory encoding and recall. Our findings are consistent with and extend findings based on transcranial magnetic stimulation suggesting a causal role of key DMN nodes during episodic memory formation (Warren KN et al. 2019). Our causal analysis also clarifies the question previously raised in fMRI studies hinting that the DMN exerts causal influences in the resting-state (Uddin LQ *et al.* 2009). We found that the DMN exerts strong causal influences on other brain networks intrinsically and surprisingly, causal outflow from the DMN to the other networks was greater during the resting-state compared to both memory encoding and recall. These findings suggest that causal influences of the DMN may not be specific to task-induced memory encoding and recall processes, and that they may also underlie internal mental processes such as recall of autobiographical information and self-monitoring as hypothesized previously (Greicius MD *et al.* 2003; Greicius MD *et al.* 2004; Buckner RL *et al.* 2008; Raichle ME 2015). The slow time-scale coherent fluctuations within DMN and faster time-scale interactions with other networks may enable the DMN to function at the apex of a neural and cognitive representational hierarchy (Margulies DS et al. 2016). Further research with denser brain-wide sampling of electrodes in multiple cortical and subcortical regions, and a wide range of experimental tasks, are necessary to further probe the broader causal role of the DMN in internal and stimulus-driven cognition.

### Conclusions

Our study advances foundational knowledge of the neural mechanisms underlying the large-scale functional organization of the human DMN. Findings provide new insights into the neurophysiological basis and intrinsic spectro-temporal organization of the DMN as well as its integrity and cross-network dynamic interactions during episodic memory formation. Findings further demonstrate that the DMN supports external stimulus-driven cognition, and not just internal mental processes (Sormaz M *et al.* 2018; Turnbull A, HT Wang, C Murphy, et al. 2019; Turnbull A, HT Wang, JW Schooler, et al. 2019). More generally, our study provides a deeper understanding of the electrophysiological mechanisms underlying operation of the DMN during cognition and its behavioral relevance. Extensions of this work with denser iEEG recordings and more fine-grained brain parcellations will further help advance knowledge of its operating mechanisms and inform the neural basis of DMN dysfunction, which impacts almost all psychiatric and neurological disorders including Alzheimer’s disease, schizophrenia, epilepsy, anxiety, depression, autism, and ADHD (Menon V 2011).

## Methods

### UPENN-RAM iEEG recordings

We examined iEEG recordings from 102 patients shared by Kahana and colleagues at the University of Pennsylvania (UPENN) (obtained from the UPENN-RAM public data release under release ID “Release_20171012”, released on 12 October, 2017) (Jacobs J *et al.* 2016). iEEG recordings were downloaded from a UPENN-RAM consortium hosted data sharing archive (UPENN-RAM). Prior to data collection, research protocols and ethical guidelines were approved by the Institutional Review Board at the participating hospitals and informed consent was obtained from the participants and guardians (Jacobs J *et al.* 2016). Details of all the recordings sessions and data pre-processing procedures are described by Kahana and colleagues (Jacobs J *et al.* 2016). Briefly, iEEG recordings were obtained using subdural grids and strips (contacts placed 10 mm apart) or depth electrodes (contacts spaced 5–10 mm apart) using recording systems at each clinical site. iEEG systems included DeltaMed XlTek (Natus), Grass Telefactor, and Nihon-Kohden EEG systems. Electrodes located in brain lesions or those which corresponded to seizure onset zones or had significant interictal spiking or had broken leads, were excluded from analysis.

Anatomical localization of electrode placement was accomplished by co-registering the postoperative computed CTs with the postoperative MRIs using FSL (FMRIB (Functional MRI of the Brain) Software Library), BET (Brain Extraction Tool), and FLIRT (FMRIB Linear Image Registration Tool) software packages. Preoperative MRIs were used when postoperative MRIs were not available. The resulting contact locations were mapped to MNI space using an indirect stereotactic technique and OsiriX Imaging Software DICOM viewer package. We used the Yeo cortical atlas (Yeo BT *et al.* 2011) in volumetric space (MNI volumetric coordinates) for mapping electrodes to the default mode, dorsal attention, ventral attention, frontoparietal, visual, motor, and limbic networks. We used the Brainnetome atlas (Fan L et al. 2016) to demarcate the hippocampus and incorporated electrodes from this region into the DMN, as the hippocampus is an important constituent subcortical node of the DMN (Greicius MD *et al.* 2003). Out of 102 individuals and 12,780 electrodes, data from 36 individuals and 879 electrodes were used for subsequent analysis based on electrode placement in these networks of interest.

iEEG signals were sampled at 1,000 Hz. The two major concerns when analyzing interactions between closely spaced intracranial electrodes are volume conduction and confounding interactions with the reference electrode (Burke JF et al. 2013). Hence bipolar referencing was used to eliminate confounding artifacts and improve the signal-to-noise ratio of the neural signals, consistent with previous studies using UPENN-RAM iEEG data (Burke JF *et al.* 2013; Ezzyat Y et al. 2018). Signals recorded at individual electrodes were converted to a bipolar montage by computing the difference in signal between adjacent electrode pairs on each strip, grid, and depth electrode and the resulting bipolar signals were treated as new virtual electrodes originating from the midpoint between each contact pair. Among the electrodes in the DMN, ~81% were depth electrodes, ~10% were grid electrodes, and ~9% were strip electrodes. For the non-DMN electrodes, these numbers were ~71%, ~17%, and ~12% respectively, indicating that the distributions of the type of electrodes in the DMN and non-DMN brain regions were similar. Line noise (60 Hz) and its harmonics were removed from the bipolar signals and finally each bipolar signal was Z-normalized by removing mean and scaling by the standard deviation. A fourth order two-way zero phase lag Butterworth filter was used in all spectral analyses.

### iEEG verbal memory encoding and recall, and resting-state conditions

Patients performed multiple trials of a “free recall” experiment, where they were presented with a list of words and subsequently asked to recall as many as possible from the original list (**Figure 1**). Details of the task are described elsewhere (Solomon EA *et al.* 2017; Solomon EA *et al.* 2019). We analyzed iEEG epochs from the encoding and recall periods of the “free recall” task similar to previous iEEG studies (Miller KJ et al. 2009; Yanagisawa T et al. 2012; Horak PC et al. 2017; Norman Y et al. 2017). Resting-state epochs consisted of intervals prior to each encoding block in which participants were asked to fixate on the screen and alerted to the upcoming task. This choice of resting-state epochs is consistent with previous iEEG and fMRI studies (Garrison KA et al. 2015; Kamp T et al. 2018; Smith V *et al.* 2018; Betzel RF et al. 2019; Das A and V Menon 2020). Moreover, in a previous study we showed that these epochs accurately reproduced findings from conventional resting-state iEEG data acquired from an independent cohort (Das A and V Menon 2020). For the resting-state epochs, we extracted 10-second iEEG recordings (epochs) prior to the beginning of each encoding block. To reduce boundary and carry over effects, we discarded 3 seconds each of iEEG data from the beginning and end of each epoch, resulting in multiple 4 second epochs. The encoding and recall epochs were 30-seconds for each trial. Data from each epoch was analyzed separately and specific measures were averaged across trials. The duration of memory encoding and recall, and resting-state epochs were matched to preclude trial-length effects.

### iEEG analysis of intra- and cross-network phase synchronization

Intra-network phase synchronization was assessed between all electrode pairs across distinct brain regions within the network. To minimize potential bias from sampling of electrodes with a specific brain region, we did not include electrode pairs within individual brain regions which were determined using the Brainnetome atlas (Fan L *et al.* 2016) as noted above. For example, for intra-DMN analysis, we computed phase synchronization between the PCC/precuneus and hippocampus electrodes but not between electrodes within the PCC/precuneus or the hippocampus. Cross-network phase synchronization was assessed between all electrode pairs across networks.

We used phase locking value (PLV) to compute phase synchronization between two time-series (Lachaux JP *et al.* 1999). We first calculated the instantaneous phases of the two signals by using the analytical signal approach based on the Hilbert transform (Bruns A 2004). Given time-series *x*(*t*), *t* = 1,2,…, *M*, its complex-valued analytical signal *z*(*t*) can be computed as

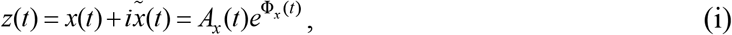

where _*i*_ denotes the square root of minus one, 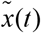 is the Hilbert transform of *x*(*t*), and *A_x_*(*t*) and Φ(*t*) are the instantaneous amplitude and instantaneous phase respectively and can be given by

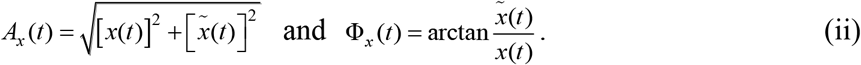

The Hilbert transform of *x*(*t*) was computed as

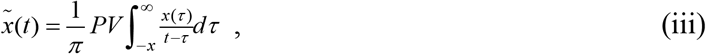

where *PV* denotes the Cauchy principal value. MATLAB function “hilbert” was used to calculate the Hilbert transform in our analysis. Given two time-series *x*(*t*) and *y*(*t*), where *t* = 1,2,…, *M*, the PLV (zero-lag) can be computed as

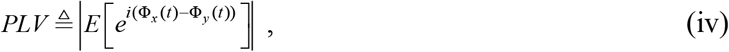

where Φ_*y*_(*t*) is the instantaneous phase for time-series *y*(*t*), |·| denotes the absolute value operator, *E* [·] denotes the expectation operator, and _*i*_ denotes the square root of minus one.

### iEEG analysis of power spectral density

To calculate average power, we first filtered the iEEG time-series in the frequency band of interest and power, after removing the linear trend, was calculated as the sum of the squares of the amplitudes of the iEEG time-series divided by the length of the time-series.

### iEEG analysis of phase transfer entropy (PTE) and causal dynamics

Causal interactions between networks was assessed using phase transfer entropy (PTE) between all electrode pairs across networks. PTE is a nonlinear measure of the directionality of information flow between time-series (Lobier M *et al.* 2014). Given two time-series {*x_i_*} and {*y_i_*}, where *i* = 1, 2,…, *M*, instantaneous phases were first extracted using the Hilbert transform. Let 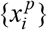 and 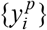, where *i* = 1,2,…, *M*, denote the corresponding phase time-series. If the uncertainty of the target signal 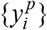 at delay *τ* is quantified using Shannon entropy, then the PTE from driver signal 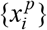 to target signal 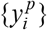 can be given by

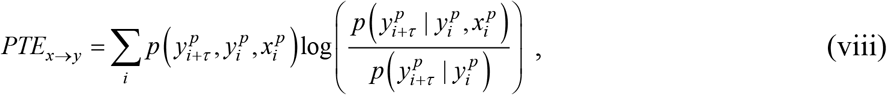

where the probabilities can be calculated by building histograms of occurences of singles, pairs, or triplets of instantaneous phase estimates from the phase time-series (Hillebrand A, P Tewarie, Ev Dellen, *et al.* 2016). For our analysis, the number of bins in the histograms was set as 3.49× *STD*×*M*^-1/3^ and delay *τ* was set as 2*M*/*M*_±_, where _*STD*_ is average standard deviation of the phase time-series 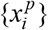 and 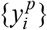 and *M*_±_ is the number of times the phase changes sign across time and channels (Hillebrand A, P Tewarie, Ev Dellen*, et al.* 2016). Note that PTE is robust against the choice of the delay *τ* and the number of bins for forming the histograms and variations in these parameters do not change the results much (Hillebrand A, P Tewarie, Ev Dellen, *et al.* 2016). For PTE estimation, we used the broadband signal rather than the filtered signal, since causality estimation is very sensitive to filtering (see (Barnett L and AK Seth 2011) for a detailed discussion on this).

### Statistical analysis

Nonparametric tests were used for statistical analysis of intra-DMN and its cross-network interactions. For phase synchronization analysis, we used the Mann-Whitney U-test, with FDR-corrections (*q*<0.05) across all frequency bands and networks to correct for multiple comparisons. For comparison of phase synchronization of memory encoding and recall, with the resting-state condition, we used the Mann-Whitney U-test with FDR-corrections (*q*<0.05) across task conditions and frequency bands. For comparison of phase synchronization values for successful and unsuccessful memory encoding and recall periods, Wilcoxon signed-rank test was used. Similarly, for phase transfer entropy and causality analysis between the DMN and the other 6 networks, we used the Wilcoxon signed-rank test. Finally, we used the Mann-Whitney U-test for control analyses of power spectral density and high-gamma band fluctuations.

## Acknowledgements

We are grateful to members of the UPENN-RAM consortia for generously sharing their unique iEEG data. We thank Drs. Paul A. Wanda, Michael V. DePalatis, Youssef Ezzyat, Richard Betzel, and Leon A. Davis for assistance with the UPENN-RAM dataset. We thank Drs. Matteo Fraschini and Arjan Hillebrand for generously sharing their MATLAB code for phase transfer entropy analysis. This research was supported by NIH grants NS086085 and EB022907.

## Supplementary Information

### I. Supplementary Methods

#### Correlation analysis of high-gamma band amplitude fluctuations

We investigated whether intra-DMN interactions showed a signature in the high-gamma band (80-160 Hz) amplitude fluctuations. iEEG signals were filtered between 80-90, 90-100,…, 150-160 Hz and the amplitude of each narrowband signal was calculated by taking the absolute value of the analytic signal obtained from the Hilbert transform (Foster BL et al. 2015). Each narrowband amplitude was then normalized with respect to the mean amplitude, i.e., expressed as fraction of the mean, to correct for 1/f decay. Normalized amplitude time-series from each band were then filtered in the slow-wave (< 4 Hz) band to generate a time series reflecting high-gamma band fluctuations in each electrode. The resulting time series was used to investigate intra- and inter-DMN interactions using Pearson correlations.

### II. Supplementary Results

#### DMN power spectral density does not differ from other networks during rest

To examine whether intra-DMN synchronization was driven by differences in the amplitude of iEEG fluctuations, we compared power spectral density (see **Methods** for details) in the DMN and the other 6 large-scale networks. This analysis revealed that power in the DMN did not differ from power in the other six networks in any of the frequency bands (all *ps*>0.05, Mann-Whitney U-test). These results suggest that synchronization within the DMN is not driven by differences in the amplitude of iEEG fluctuations.

#### DMN power spectral density does not differ from other networks during memory processing

Similar to our analysis of resting-state iEEG, we compared power spectral density in the DMN and the other 6 networks during memory encoding, and separately during memory recall. This analysis revealed that power in the DMN did not differ from power in the other six networks in any of the frequency bands (*ps*>0.05, Mann-Whitney U-test) in either the encoding or recall phases of the episodic memory task.

#### Spectral power density during memory processing does not differ from rest

We next compared the power spectral density during memory encoding or recall with power spectral density during resting-state. As with previous analyses, we randomly selected epochs from the memory encoding or recall periods to match their duration to those from the resting-state epochs, thereby ensuring that differences in network dynamics could not be explained by the differences in the duration of the epochs. For the DMN, power during memory encoding or recall did not differ from power during rest in any of the frequency bands (*ps*>0.05 in all frequency bands, Mann-Whitney U-test). This indicates that the increased phase synchronization during memory encoding/recall is not driven by differences in the amplitude of iEEG fluctuations.

#### Spectral power density during successful memory encoding/recall does not differ from unsuccessful memory encoding/recall

We compared the power spectral density during successful memory encoding/recall with the power spectral density during unsuccessful memory encoding/recall in the slow-wave (< 4 Hz) frequency band for the DMN. This analysis revealed that power during successful memory encoding/recall did not differ from power during unsuccessful memory encoding/recall (*ps*>0.05, Wilcoxon signed-rank test). This indicates that the increased intra-DMN phase synchronization during successful encoding/recall compared to unsuccessful encoding/recall in the slow-wave frequency band is not driven by differences in the amplitude of iEEG fluctuations.

#### Intra- and inter-DMN correlation in high-gamma band amplitude fluctuations did not differ from each other across conditions

Previous studies have suggested that low-frequency fluctuations in the high-gamma band (80-160 Hz) are correlated with fMRI BOLD signals (Leopold DA et al. 2003; Mantini D et al. 2007; Schölvinck ML et al. 2010; Hutchison RM et al. 2015; Lakatos P et al. 2019). Our analysis (**Supplementary Methods**) revealed that intra- and inter-DMN correlation in high-gamma band fluctuations did not differ from each other in the resting-state, memory encoding, and memory recall conditions (all *ps*>0.05, Mann-Whitney U-test).

### III. Supplementary Figures

**Figure S1.**
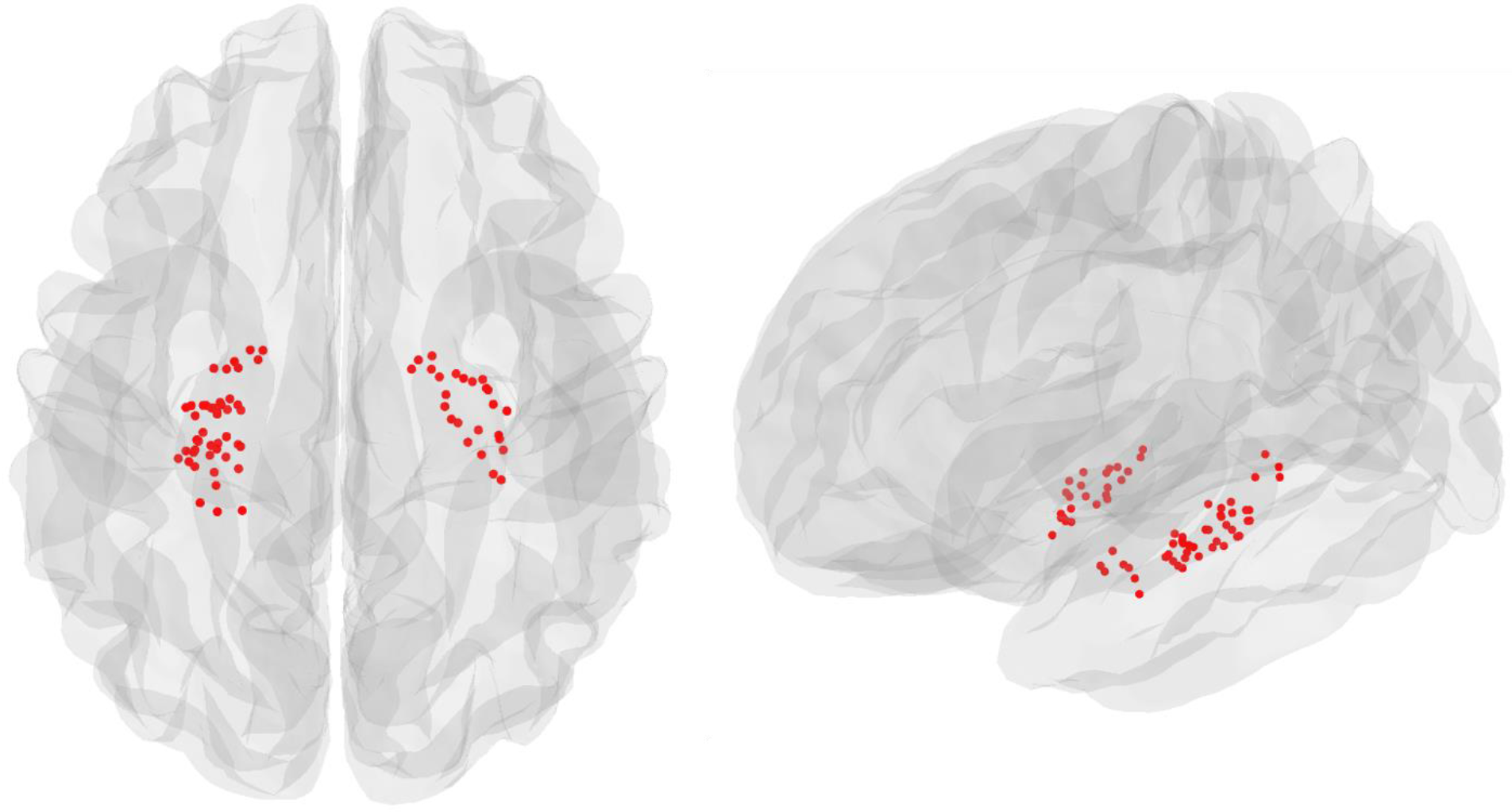
**iEEG recordings sites in the hippocampus.** Left and right hemisphere hippocampal regions were determined using the Brainnetome atlas (Fan L et al. 2016).

**Figure S2. Movie showing the Yeo cortical atlas (Yeo BT et al. 2011) (in yellow) and the iEEG recordings sites in the DMN (in red).** Note: Click on the link or copy and paste the link on a web browser below to run the movie.

https://raw.githubusercontent.com/anupdas777/Yeo-DMN-movie/master/yeo_elec.mov

**Figure S3.**
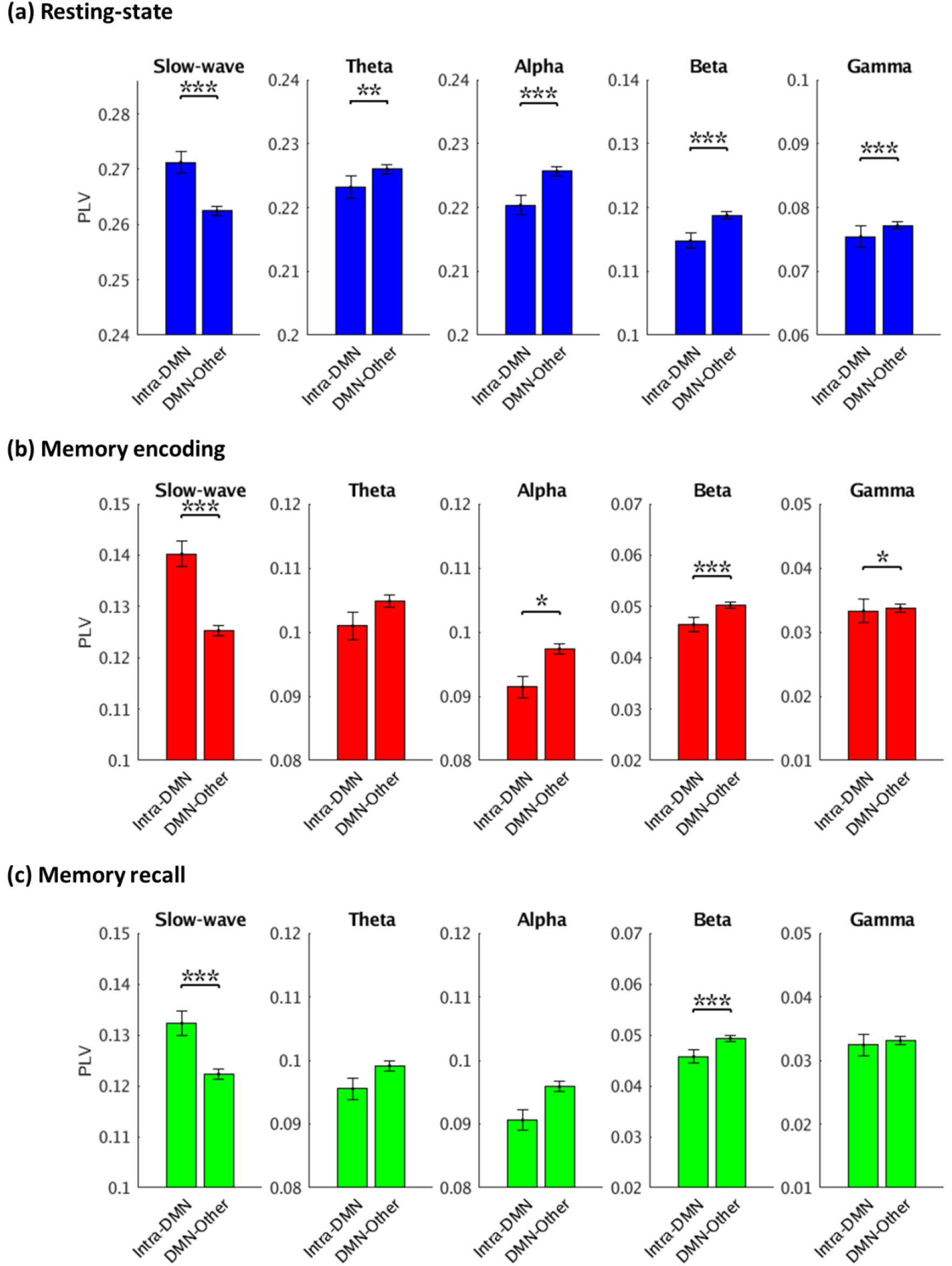
**Intra-DMN phase synchronization when hippocampus electrodes were excluded from the DMN during (a) Resting-state, (b) Memory encoding, and (c) Memory recall conditions.** Error bars denote standard error of the mean (SEM) across all pairs of electrodes. *** *p* < 0.001, ** *p* < 0.01, \**p* < 0.05 (FDR-corrected *q*<0.05).

**Figure S4.**
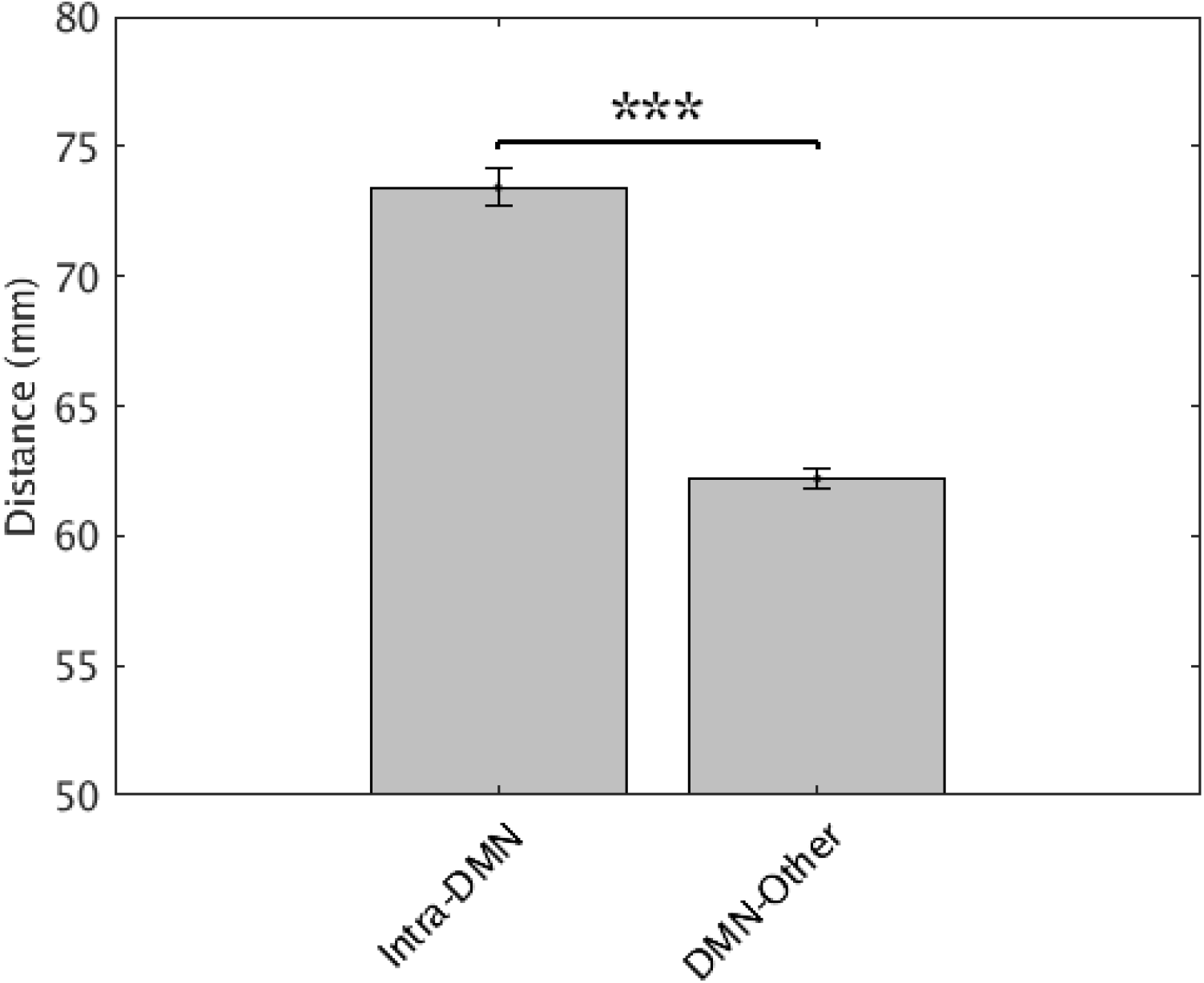
**Intra-DMN electrodes distance, compared to distance of DMN electrodes from electrodes in the 6 other brain networks.** Intra-DMN electrode distance was higher than the DMN electrodes distance with other networks (DMN-Other). Error bars denote standard error of the mean (SEM) across all pairs of electrodes. *** *p* < 0.001.

**Figure S5.**
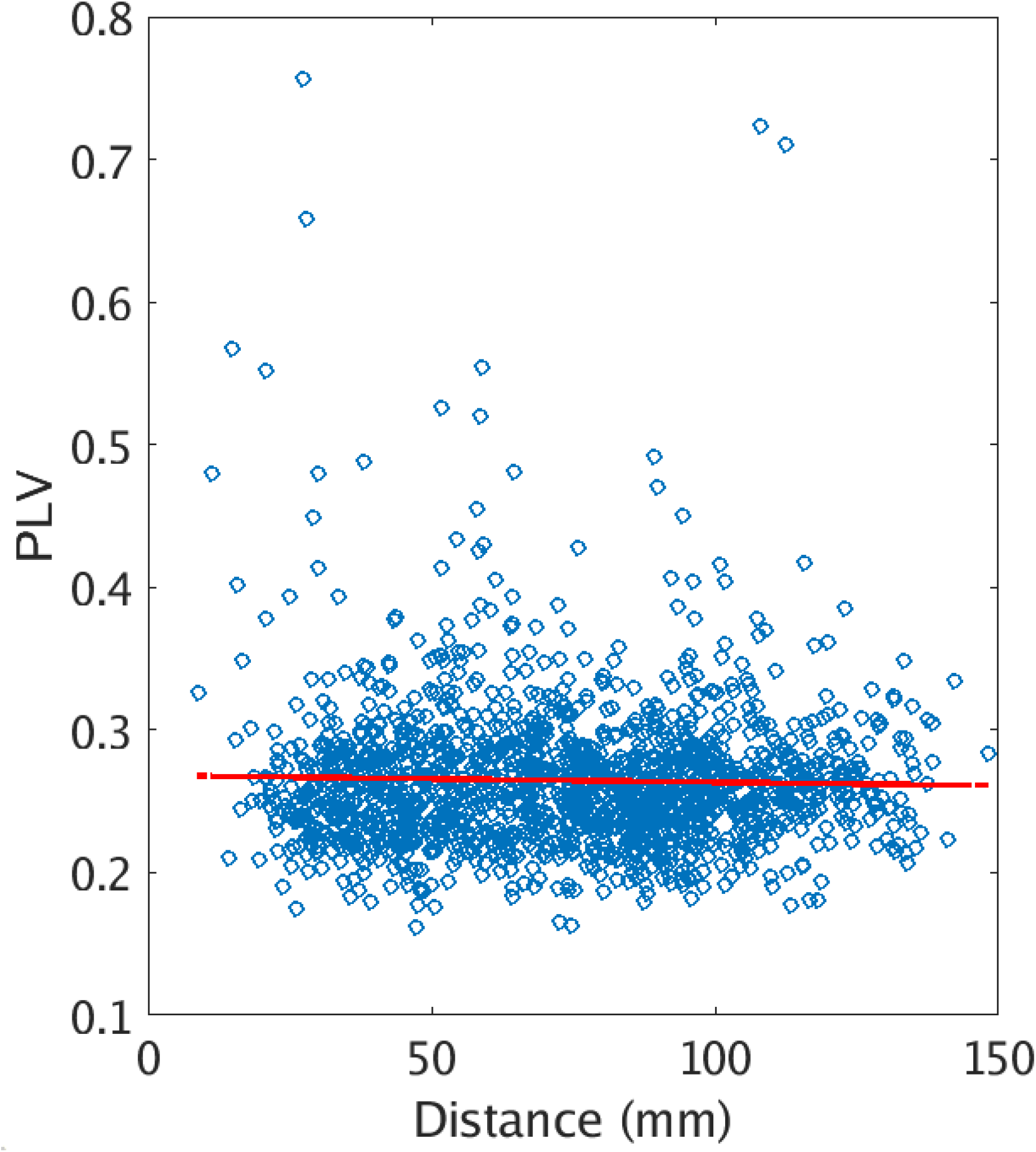
**Relationship between phase locking value (PLV) and inter-electrode distance in resting-state iEEG.** Correlation between PLV and inter-electrode distance in the DMN was not significant (*p* > 0.27).

**Figure S6.**
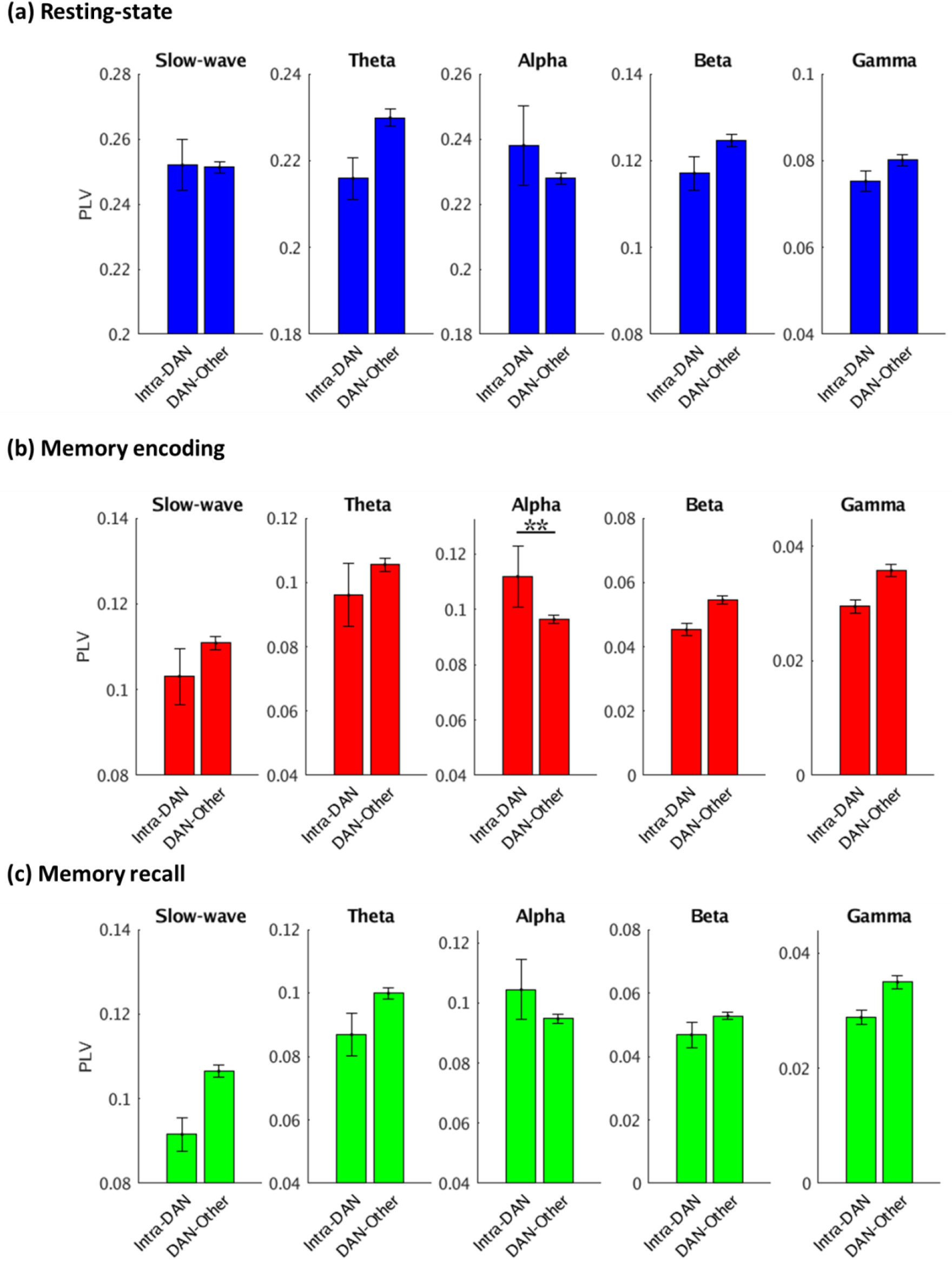
**Intra-DAN (DAN: dorsal attention network) phase synchronization with 6 other networks during (a) Resting-state, (b) Memory encoding, and (c) Memory recall conditions.** Error bars denote standard error of the mean (SEM) across all pairs of electrodes. ** *p* < 0.01 (FDR-corrected *q*<0.05).

**Figure S7.**
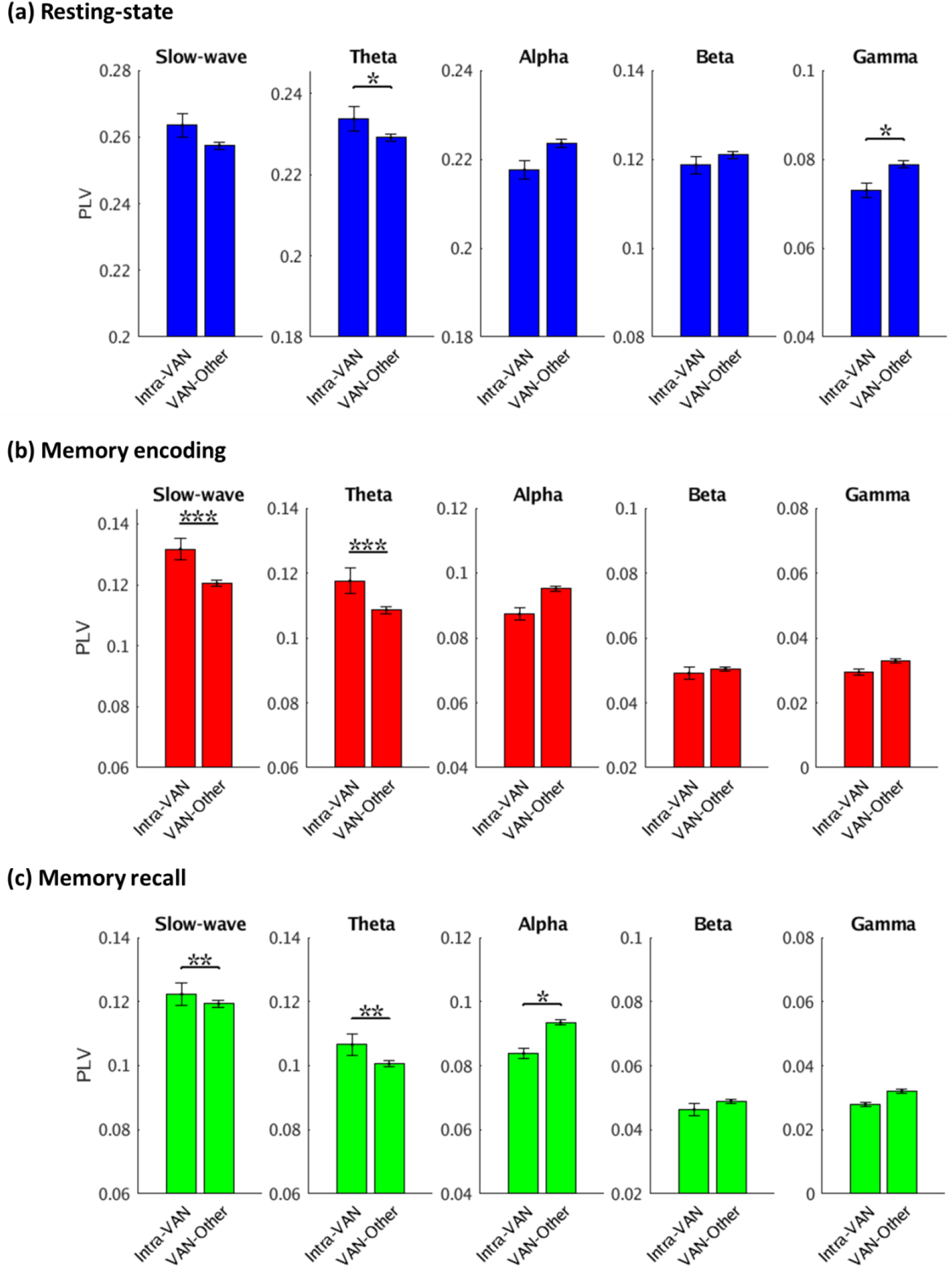
**Intra-VAN (VAN: Ventral attention network) phase synchronization with other 6 networks during (a) Resting-state, (b) Memory encoding, and (c) Memory recall conditions.** Error bars denote standard error of the mean (SEM) across all pairs of electrodes. *** *p* < 0.001, ** *p* < 0.01, \**p* < 0.05 (FDR-corrected *q*<0.05).

**Figure S8.**
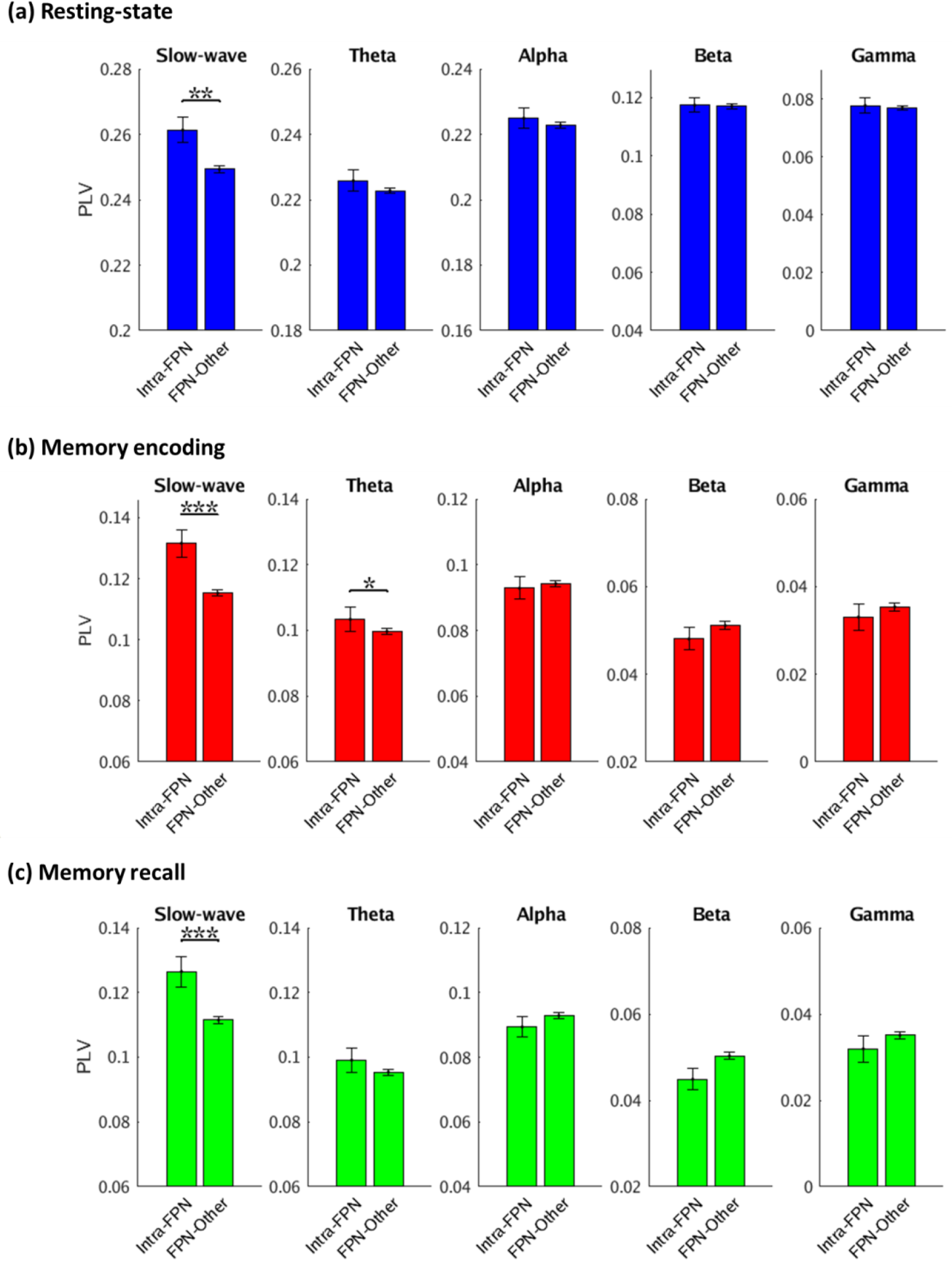
**Intra-FPN (FPN: Frontoparietal Network) phase synchronization with 6 other networks during (a) Resting-state, (b) Memory encoding, and (c) Memory recall conditions.** Error bars denote standard error of the mean (SEM) across all pairs of electrodes. *** *p* < 0.001, ** *p* < 0.01, * *p* < 0.05 (FDR-corrected *q*<0.05).

**Figure S9.**
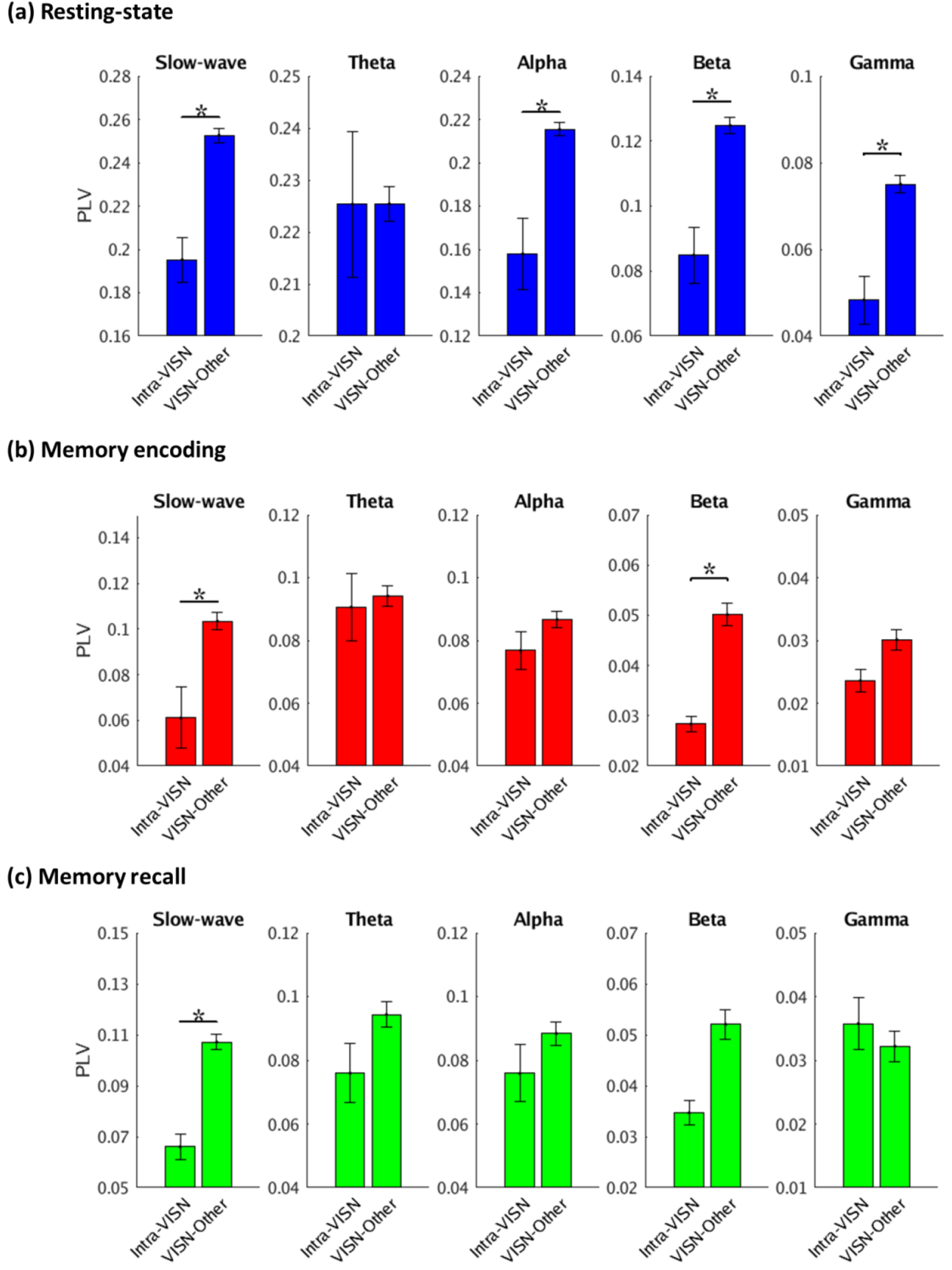
**Intra-VISN (VISN: Visual Network) phase synchronization with 6 other networks during (a) Resting-state, (b) Memory encoding, and (c) Memory recall conditions.** Error bars denote standard error of the mean (SEM) across all pairs of electrodes. * *p* < 0.05 (FDR-corrected *q*<0.05).

**Figure S10.**
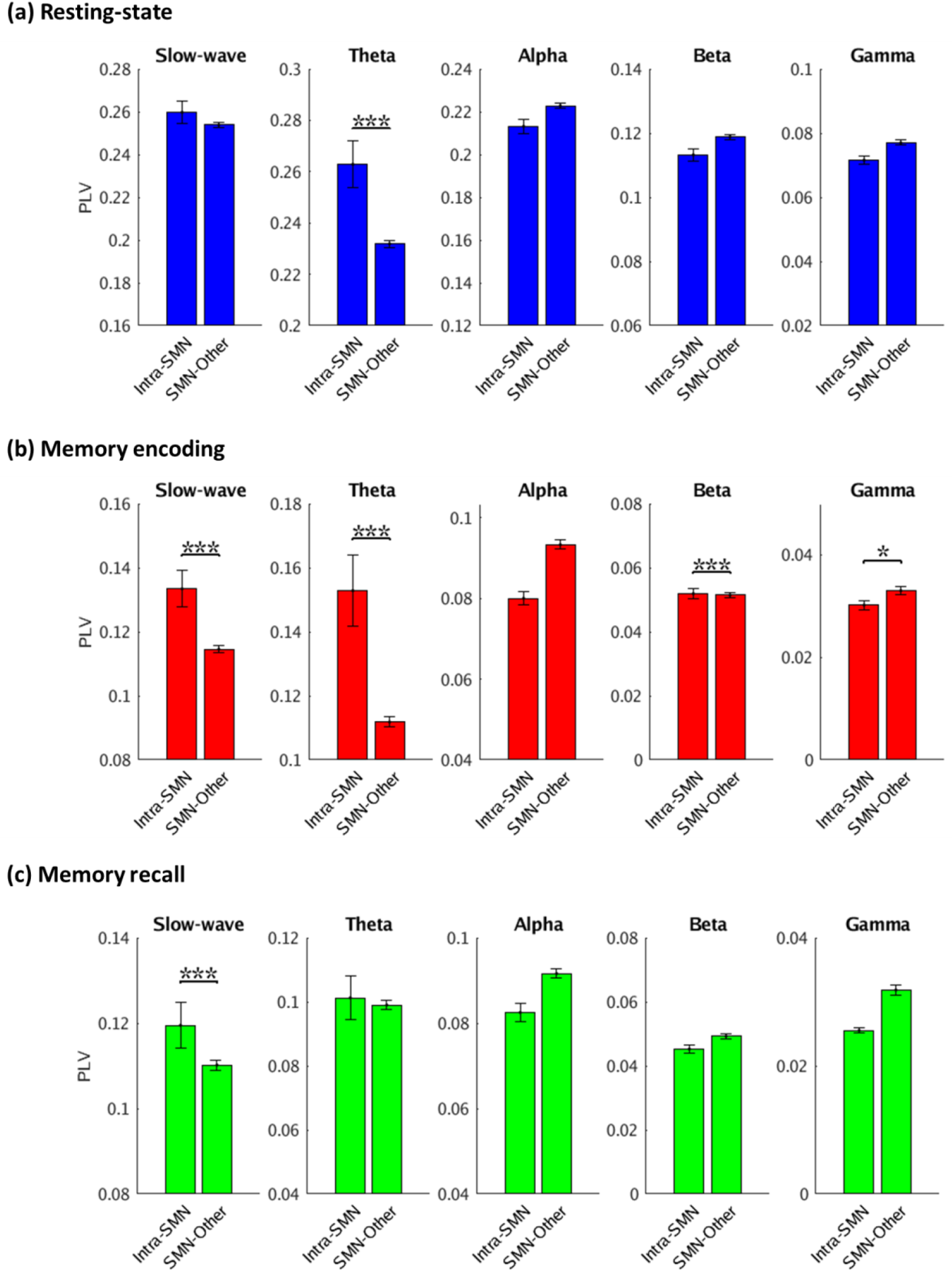
**Intra-SMN (SMN: Somatomotor Network) phase synchronization with 6 other networks during (a) Resting-state, (b) Memory encoding, and (c) Memory recall conditions.** Error bars denote standard error of the mean (SEM) across all pairs of electrodes. *** *p* < 0.001, * *p* < 0.05 (FDR-corrected *q*<0.05).

**Figure S11.**
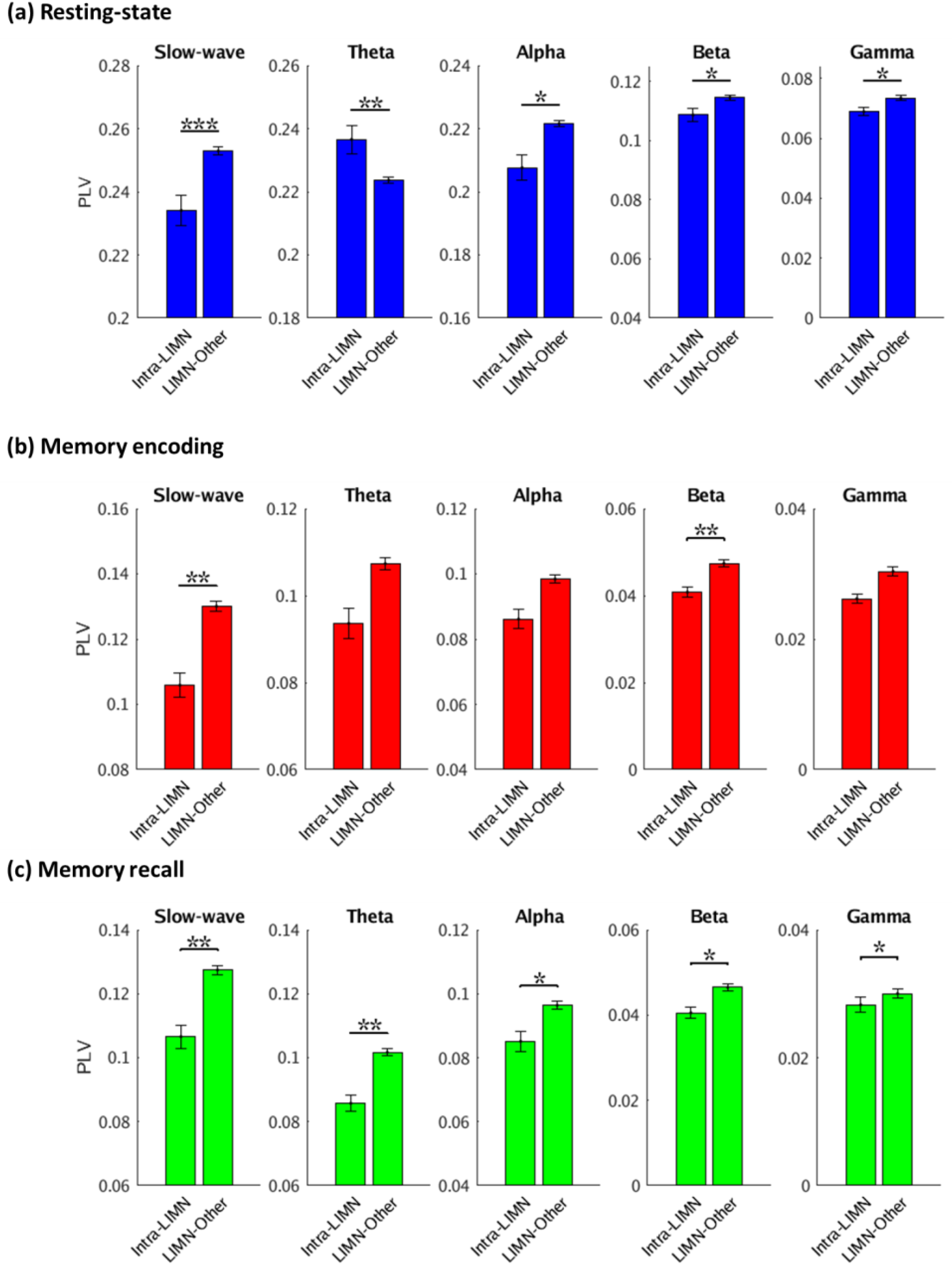
**Intra-LIMN (LIMN: Limbic Network) phase synchronization with 6 other networks during (a) Resting-state, (b) Memory encoding, and (c) Memory recall conditions.** Error bars denote standard error of the mean (SEM) across all pairs of electrodes. *** *p* < 0.001, ** *p* < 0.01, * *p* < 0.05 (FDR-corrected *q*<0.05).

### IV. Supplementary Tables

**Table S1.** Comparison with prior iEEG studies examining inter-regional connectivity of individual nodes of the DMN.

| Reference | Number of subjects | Brain regions/networks examined | Experimental conditions | Measures used | Major findings | Intra- vs. inter-DMN connectivity analyzed? |
| --- | --- | --- | --- | --- | --- | --- |
| Foster et al. (Foster BL et al. 2015) | 3 | Retrosplenial cortex (RSC), posterior cingulate cortex (PCC), and angular gyrus (AG) regions of the DMN | Resting-state, sleep (stages 2/3), and autobiographical episodic retrieval | Linear correlation of high-frequency band (HFB) power time-series | RSC/PCC and AG are correlated during episodic memory retrieval. | No |
| Foster et al. (Foster BL et al. 2013) | 4 | Medial temporal lobe (MTL) and retrosplenial cortex (RSC) regions of the DMN | Resting-state, mental arithmetic, and autobiographical memory retrieval | Phase synchronization | MTL-RSC theta frequency band phase locking during autobiographical retrieval. | No |
| Kucyi et al. (Kucyi A et al. 2018) | 5 | PCC/precuneus and medial prefrontal cortex (mPFC) regions of the DMN | Resting wakefulness and sleep (stages 2/3) | Linear correlation of high-frequency band (HFB) power time-series | Low-frequency ECoG activity is correlated with spontaneous BOLD fluctuations. | No |
| Kam et al. (Kam JWY et al. 2019) | 12 | Lateral temporal cortex node of the DMN, and FPN | Auditory attention task | Linear correlation | Increased theta frequency band connectivity between DMN and FPN during internally directed attention compared to externally directed attention. | No |
| Hacker et al.<br>(Hacker CD et al. 2017) | 6 | Superior frontal gyrus (SFG), DAN, FPN, SMN | Resting-state | Linear correlation | Electrocorticogram -fMRI correspondence is stronger in the theta frequency band in the DMN and FPN and alpha frequency band in the DAN and SMN. | No |
| Kucyi et al.<br>(Kucyi A et al. 2020) | 31 | Posteromedial cortex (PMC), medial prefrontal cortex (mPFC), and angular gyrus (AG) regions of the DMN, DAN | Resting-state and attention task (Gradual-onset Continuous Performance Task) | High-frequency band (HFB) power time-series | DMN-DAN HFB coupling is associated with greater sustained attention performance. | No |
| Present study | 36 | vmPFC, alPFC, STS/MTG, IPL, PCC/precuneus, and hippocampus regions of the DMN, along with DAN, VAN, FPN, VISN, SMN, LIMN | Resting-state, memory encoding, and memory retrieval | Nonlinear Phase synchronization, Nonlinear causal dynamics |  | Yes |

**Table S2.** Participant demographic information.

| **Participant ID** | **Gender** | **Age** |
| --- | --- | --- |
| 184 | M | 42 |
| 185 | M | 20 |
| 186 | M | 27 |
| 187 | F | 51 |
| 189 | M | 22 |
| 193 | M | 37 |
| 195 | M | 44 |
| 196 | M | 18 |
| 200 | M | 25 |
| 202 | F | 29 |
| 203 | F | 36 |
| 204 | F | 25 |
| 207 | F | 39 |
| 215 | F | 50 |
| 222 | F | 20 |
| 223 | F | 42 |
| 228 | F | 58 |
| 229 | M | 28 |
| 230 | F | 56 |
| 231 | M | 20 |
| 232 | M | 27 |
| 234 | M | 25 |
| 247 | F | 61 |
| 251 | M | 31 |
| 260 | F | 57 |
| 264 | F | 52 |
| 268 | F | 32 |
| 274 | F | 44 |
| 275 | M | 41 |
| 283 | F | 29 |
| 286 | F | 57 |
| 292 | F | 39 |
| 297 | M | 24 |
| 298 | F | 24 |
| 299 | M | 43 |
| 304 | F | 33 |
| 310 | M | 20 |

**Table S3.** Number of electrodes in each brain network. DMN: default mode network, DAN: dorsal attention network, VAN: ventral attention network, FPN: frontoparietal network, VISN: visual network, SMN: somatomotor network, LIMN: limbic network.

| Brain network | Number of electrodes* (n) | Number of participants | Participant IDs |
| --- | --- | --- | --- |
| DMN | 347 | 33 | 185, 186, 187, 193, 195, 196, 200, 203, 204, 207, 215, 222, 223, 228, 229, 230, 231, 232, 234, 247, 260, 264, 268, 274, 275, 283, 286, 292, 297, 298, 299, 304, 310 |
| DAN | 52 | 24 | 184, 185, 186, 187, 189, 193, 195, 196, 200, 202, 204, 222, 223, 230, 234, 260, 264, 275, 283, 286, 297, 298, 299, 310 |
| VAN | 159 | 30 | 184, 185, 186, 189, 193, 195, 196, 200, 203, 204, 207, 222, 223, 228, 230, 231, 232, 247, 251, 260, 264, 274, 275, 283, 286, 292, 297, 298, 299, 310 |
| FPN | 123 | 30 | 184, 185, 186, 189, 193, 195, 196, 200, 203, 204, 207, 222, 223, 228, 230, 232, 234, 247, 260, 264, 268, 274, 275, 283, 286, 297, 298, 299, 304, 310 |
| VISN | 19 | 8 | 185, 187, 202, 234, 264, 268, 283, 297 |
| SMN | 97 | 23 | 184, 186, 189, 193, 195, 200, 203, 204, 207, 223, 230, 231, 232, 251, 260, 268, 274, 275, 283, 292, 297, 299, 310 |
| LIMN | 82 | 21 | 185, 186, 196, 200, 203, 204, 207, 223, 230, 234, 247, 260, 264, 268, 274, 275, 283, 297, 298, 299, 304 |

\*The encoding session file for subject 185 was missing. For the memory encoding task, the number of electrodes (n) were 345, 51, 158, 119, 16, and 81 for DMN, DAN, VAN, FPN, VISN, and LIMN respectively.

**Table S4.**
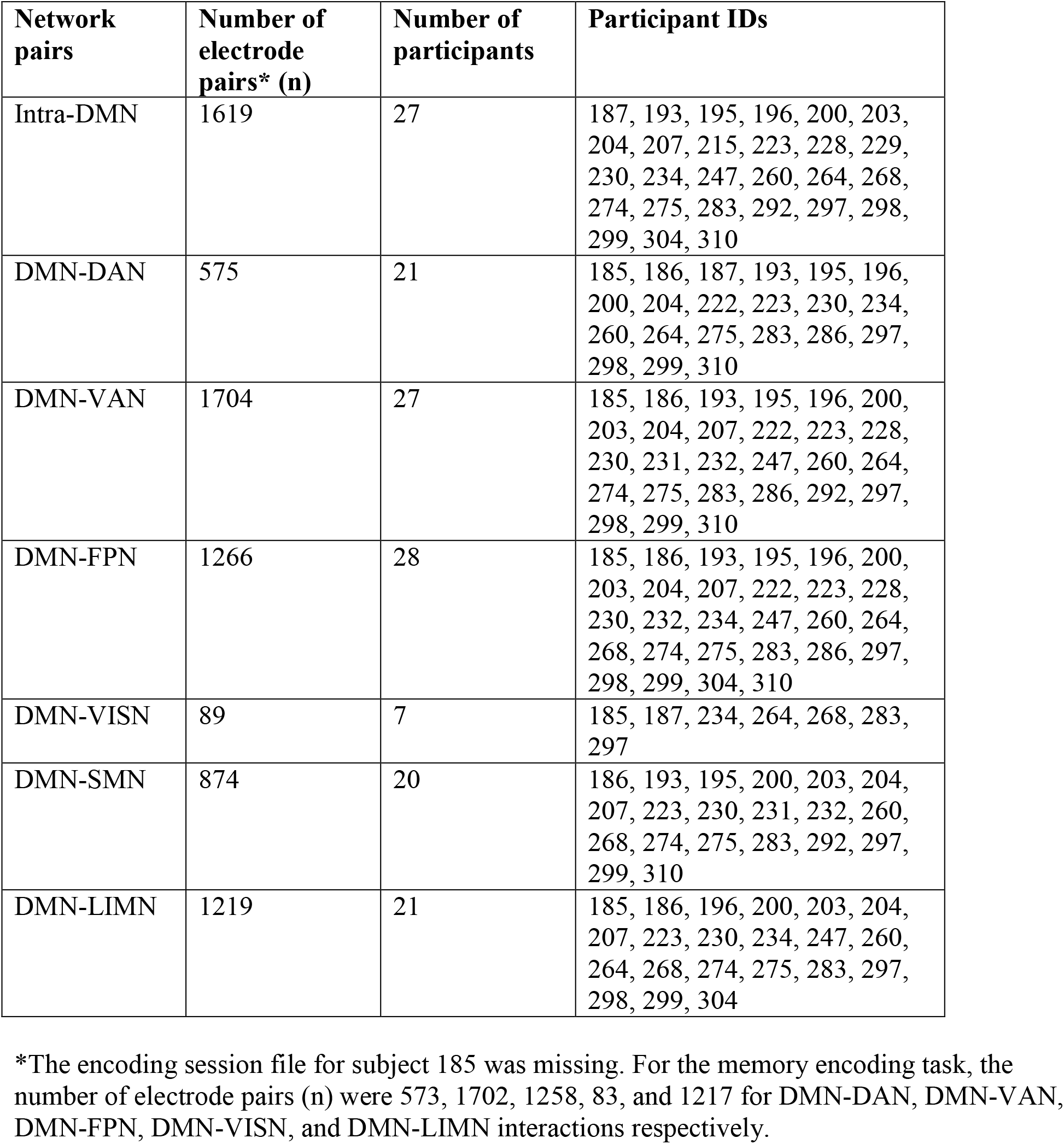
Number of electrode pairs used in intra- and cross-network connectivity analysis. DMN: default mode network, DAN: dorsal attention network, VAN: ventral attention network, FPN: frontoparietal network, VISN: visual network, SMN: somatomotor network, LIMN: limbic network.

**Table S5.** Number of electrodes in different nodes of the DMN. vmPFC: ventromedial prefrontal cortex, alPFC: anterior lateral prefrontal cortex, STS: superior temporal sulcus, MTG: middle temporal gyrus, IPL: inferior parietal lobule, PCC: posterior cingulate cortex.

| **DMN node** | **Number of electrodes** |
| --- | --- |
| vmPFC | 30 |
| alPFC | 82 |
| STS/MTG | 135 |
| IPL | 21 |
| PCC/precuneus | 9 |
| Hippocampus | 70 |

**Table S6.** Summary of intra- and inter-network PLVs across frequency bands and conditions for different networks. DMN: default mode network, DAN: dorsal attention network, VAN: ventral attention network, FPN: frontoparietal network, VISN: visual network, SMN: somatomotor network, LIMN: limbic network.

| **Brain network** | **Whether intra-network PLV > inter-network PLV in slow-wave frequency band (< 4 Hz) across conditions (Yes/No)** | **Whether inter-network PLV > intra-network PLV in higher frequency bands (theta, alpha, beta, and gamma) across conditions (Yes/No)** |
| --- | --- | --- |
| DMN | Yes | Yes |
| DAN | No | No |
| VAN | No | No |
| FPN | Yes | No |
| VISN | No | No |
| SMN | No | No |
| LIMN | No | No |

